# Towards middle-up analysis of polyclonal antibodies: subclass-specific *N*-glycosylation profiling of murine immunoglobulin G (IgG) by means of HPLC-MS

**DOI:** 10.1101/2020.07.16.205930

**Authors:** Constantin Blöchl, Christof Regl, Christian G. Huber, Petra Winter, Richard Weiss, Therese Wohlschlager

## Abstract

Advanced analytical strategies including top-down and middle-up HPLC-MS approaches have become powerful alternatives to classical bottom-up analysis for the characterization of therapeutic monoclonal antibodies. Here, we assess feasibility of middle-up analysis of polyclonal IgGs posing additional challenges due to extensive sequence variability. The presented workflow is based on Fc/2 portions as conserved subunits of IgGs and enables global profiling of subclasses and their glycosylation patterns, both of which influence IgG effector functions. To obtain subunits of murine IgGs, we established digestion with the bacterial protease SpeB. The resulting Fc/2 portions characteristic of different subclasses were subsequently analysed by ion-pair reversed-phase HPLC hyphenated to high-resolution mass spectrometry allowing relative quantification of IgG subclasses and their *N*-glycosylation variants. In order to assess method capabilities in an immunological context, we applied the analytical workflow to polyclonal antibodies obtained from BALB/c mice immunized with the grass pollen allergen Phl p 6. This analysis simultaneously revealed a shift in IgG subclasses and Fc-glycosylation patterns in total and antigen-specific IgGs from different mouse cohorts. Eventually, Fc/2 characterization may reveal other protein modifications including oxidation, amino acid exchanges, and C-terminal lysine as demonstrated for monoclonal IgGs, which may be implemented for quality control of functional antibodies.

## Introduction

Emergence of novel analytical technologies has driven the development of intact protein and subunit analysis approaches, providing information on protein integrity and coexistence of protein variants.^1^ These advanced strategies are especially attractive for the characterization of monoclonal antibodies (mAbs), i.e. purified therapeutic proteins of a single defined amino acid sequence. In this context, intact protein analysis reveals protein heterogeneity arising from glycosylation and other post-translational modifications (PTMs) as has been demonstrated for several therapeutic mAbs and an Fc-fusion protein.^2-8^ Alternative strategies for mAb characterization are based on protein subunits obtained by reduction of disulphide bonds or limited proteolysis.^9-11^ Subsequent mass determination of protein subunits is referred to as “middle-up”, additional gas phase fragmentation and fragment mass determination as “middle-down” analysis.^12^

In contrast to monoclonal immunoglobulin G (IgG)-type antibodies, native polyclonal IgGs occurring in biological samples, e.g. serum, exhibit a near infinite number of sequence variants, which arise from their antigen-specific variable regions. IgGs generally consist of an antigen binding fragment (Fab) determining antigen specificity and a crystallisable fragment (Fc) conveying effector functions to the antibody. Based on the conserved amino acid sequences of the Fc domain, IgGs are divided into subclasses, i.e. IgG1 to 4 in humans, which differ in their binding to Fc receptors and hence their effector functions. Structure and function of all IgG subclasses is additionally tuned by *N*-glycosylation of the Fc domain.^13^ Substitution of the conserved *N*-glycosylation site with extended *N*-glycans, for example, favours an open conformation of the Fc domain, while truncation of *N*-glycan structures or complete deglycosylation of the IgG induce a more closed state.^14,15^ These structural alterations modulate IgG-mediated effector functions, e.g. antibody-dependent cellular cytotoxicity (ADCC), complement-dependent cytotoxicity (CDC), and anti-inflammatory responses.^16,17^ In addition to these structure-function relationships, IgG glycosylation profiles have been shown to be associated with the physiological state of an organism.^18,19^ For example, changes in IgG glycosylation have been observed for various indications, including autoimmune diseases and cancer, both in humans^20-22^ and in mouse models^23^ of human disease. The effect of antigen stimulation on IgG glycosylation patterns as part of the immune response has been studied in mice.^24^

As antibody function is controlled via changes in both antibody subclass selection and antibody glycosylation^25,26^, monitoring of IgG glycosylation in a subclass-specific manner may reveal fine-tuning of effector functions and provide new insights into immune regulatory processes. Analysis of polyclonal IgGs with respect to subclass abundances and glycosylation patterns is conventionally based on ELISA and glycopeptides/released glycans, respectively.^24,27-29^ Specifically, ELISA provides comparative values for a certain IgG subclass in multiple samples. Differences in the affinity of secondary antibodies, however, preclude direct comparison of different IgG subclasses within a sample. With respect to glycosylation of polyclonal antibodies, site-specific information may be derived from glycopeptide data facilitating assignment of certain IgG subclasses.^27,30-32^ Assessment of glycoform heterogeneity at the intact protein level as performed for mAbs, however, is precluded by the vast number of sequence variants. Specific proteolytic cleavage in the hinge may allow segregation of the variable Fab domain from the Fc portion to facilitate middle-up/middle-down analysis of this conserved domain. As determinant for IgG subclasses, characterization of Fc domains may thus enable subclass-specific Fc glycosylation monitoring of polyclonal IgGs.

Here, we explore the potential of middle-up/middle-down analysis for polyclonal IgGs from mouse representing the main model organism for immunological studies. In order to obtain Fc/2 subunits, we established proteolytic digestion of murine IgGs with SpeB, a bacterial protease of previously unknown sequence specificity. Indeed, all murine IgG subclasses and isotypes (1/1i, 2a/c, 2b/2bi, and 3) were amenable to SpeB proteolysis. HPLC-MS analysis of the obtained Fc/2 subunits enabled assignment of IgG subclasses and isotypes in a middle-down approach. Middle-up analysis simultaneously provided qualitative and quantitative information on IgG glycosylation in a subclass-specific fashion. Finally, we demonstrate capabilities of the described workflow in the analysis of polyclonal IgGs from serum of BALB/c mice immunized with timothy grass pollen allergen Phl p 6.^33^ Our approach provides global information on IgG subclass abundances and their respective glycoforms and paves the way towards the characterization of polyclonal IgG proteoforms.

## Results and Discussion

### Analytical strategy

Inspired by recent advances in the characterization of therapeutic mAbs at the subunit level, we set out to establish a middle-up workflow for Fc-glycosylation analysis in polyclonal murine IgGs.^10,34^ Specifically, we aimed at the analysis of Fc/2 as the determinant for IgG subclasses (see Fig. 1). Subunits of human antibodies may be obtained upon digestion with the IdeS protease, which is a well established tool for characterization of therapeutic monoclonal IgGs. With respect to polyclonal human IgGs, digestion with IdeS was used to generate Fc/2 for the determination of IgG1 and IgG2 allotypes^35^ and for monitoring of oxidation in intravenous immunoglobulin preparations (IVIG).^36^ While IdeS cleaves human IgGs of all subclasses, the enzyme does not act on murine IgG1 and IgG2b isoforms. We therefore assessed applicability of the bacterial protease SpeB to cleave polyclonal mouse IgGs of all subclasses within the hinge region (Fig. 1a, b). Digestion with SpeB yielded three species of IgG subunits, all approximately 25 kDa in size: light chain (LC), N-terminal part (Fd’) and C-terminal part (Fc/2) of the heavy chain (HC). Variable regions are located in LC and Fd’, whereas sequence variability of the glycosylated Fc/2 subunit is limited. In fact, each IgG subclass comprises a characteristic Fc/2 portion defined by a conserved amino acid sequence. Taking advantage of these well-defined sequence variants, we apply IP-RP-HPLC to separate Fc/2 subunits of different IgG subclasses (Fig. 1c). Noteworthy, LC and Fd’ portions do not interfere with the analysis, as they are chromatographically separated from Fc/2 subunits. Hyphenation to high-resolution Orbitrap™mass spectrometry provides qualitative and quantitative information on Fc/2 glycosylation variants (Fig. 1d).

**Figure 1.**
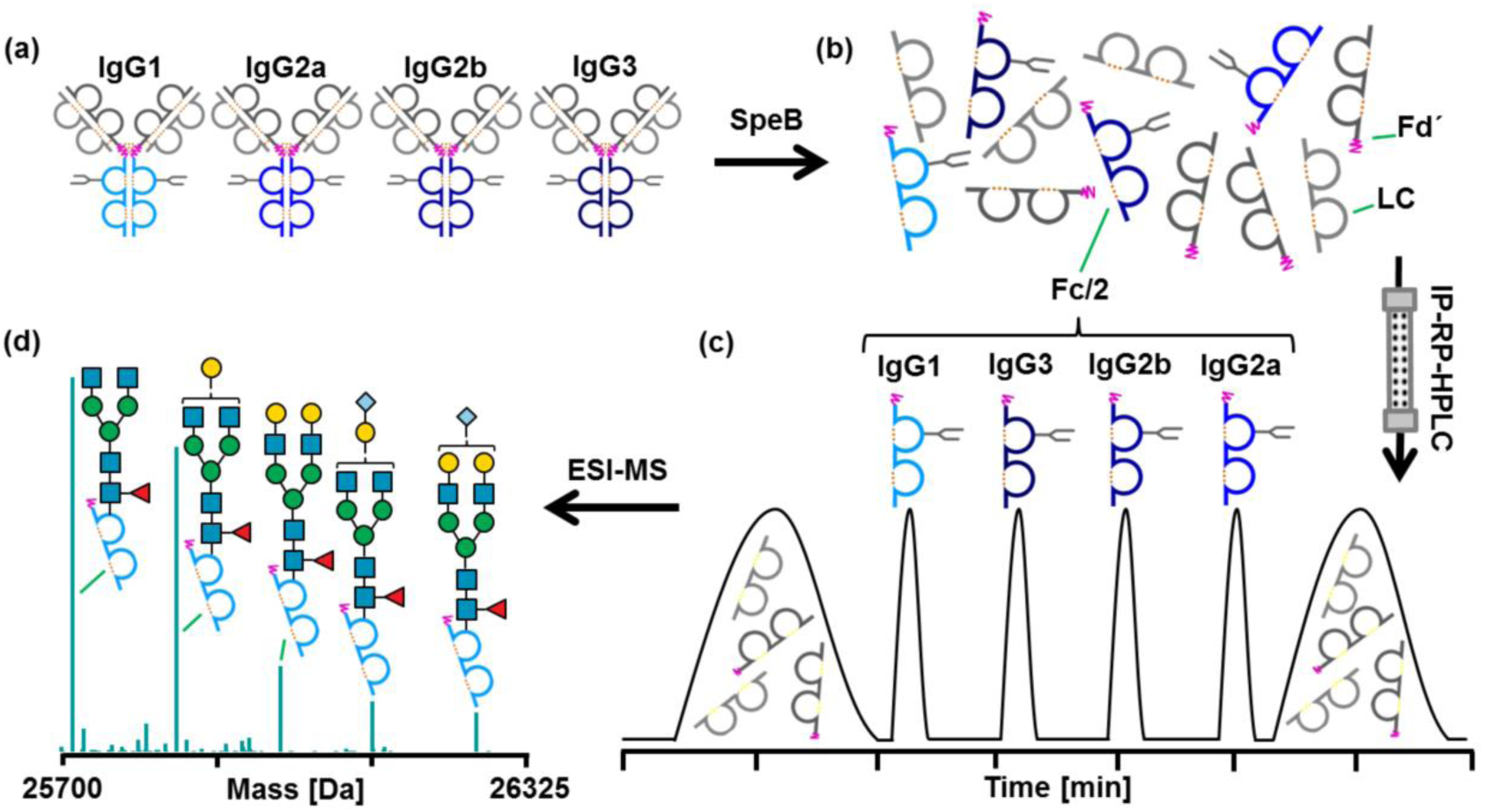
Schematic workflow for Fc/2 subunit characterization of polyclonal murine IgGs. (a) A mixture of affinity-purified polyclonal murine IgGs of different subclasses is digested with SpeB under reducing conditions. (b) Proteolytic cleavage in the hinge region (pink) generates LC (light grey), Fd’ (dark grey) and Fc/2 (blue) subunits. (c) IP-RP-HPLC of IgG subunits facilitates separation of Fc/2 portions in a subclass-specific way. (d) Glycoforms are identified for IgG subclasses based on mass spectra of Fc/2.

### Determination of SpeB cleavage products in murine IgG subclasses

In order to obtain Fc/2 subunits of all IgG subclasses described in mice, we explored the proteolytic properties of the cysteine protease SpeB, the major virulence factor of the human pathogen *Streptococcus pyogenes*. SpeB has been shown to cleave streptococcal proteins as well as human host proteins, including IgGs.^37-39^ While rather loose sequence specificity was described for SpeB, cleavage is thought to be largely determined by site accessibility and hence the three-dimensional structure of the protein substrate.^39^ Accordingly, SpeB preferentially cleaves human IgGs within the exposed hinge region, thereby generating free Fc/2 subunits.^38^ As the cleavage sites in murine IgGs were unknown, we initially set out to identify the proteolytic products resulting from SpeB cleavage of HC portions from different IgG subclasses. For this purpose we used monoclonal mouse IgGs, i.e. IgG1, IgG2a, IgG2b and IgG3, as well as polyclonal IgGs of two commonly used inbred mouse strains, BALB/c and C57BL/6. The former mouse strain comprises IgG1, IgG2a, IgG2b and IgG3 subclasses, while C57BL/6 mice feature IgG1i, IgG2c, IgG2bi and IgG3, respectively.^40^ Employing protein G affinity chromatography, we obtained mixtures of polyclonal IgGs from the serum of mouse individuals of these strains. Monoclonal antibodies as well as polyclonal IgGs were digested with SpeB for 3 h at 37 °C in the presence of L-cysteine. The generated subunits were then analysed by IP-RP-HPLC-MS, applying full MS/all ion fragmentation (AIF) mode (Supplementary Fig. S-1). HC cleavage products were subsequently determined as follows: the amino acid sequence of the HC was identified by matching y-ions (C-terminal fragments) to a candidate list of Uniprot entries; the cleavage site was localised in the identified sequence based on the experimental mass of the cleavage product; the amino acid sequence of the assigned cleavage product was verified by b-ions (N-terminal fragments). Employing this combination of middle-up and middle-down analysis, IgG subclasses from C57BL/6 could be associated with the following accession numbers: IgG1i (A0A075B5P4), IgG2bi (A0A075B5P3) and IgG2c (A0A0A6YY53). In BALB/c mice, IgG1 (P01868), IgG2a (P01863) and IgG2b (P01867) were assigned (Table 1, Supplementary Fig. S-2). IgG1 from this mouse strain corresponded to a sequence variant of the P01868 Uniprot entry comprising two amino acid exchanges (both N to D). This previously described variant was additionally confirmed by peptide analysis (Supplementary Fig. S-3).^41^ IgG3 was not detected in SpeB digests of polyclonal antibodies, which may be due to the low abundance of this subclass in serum or due to protein precipitation upon formation of cryoglobulins.^42,43^

**Table 1.**
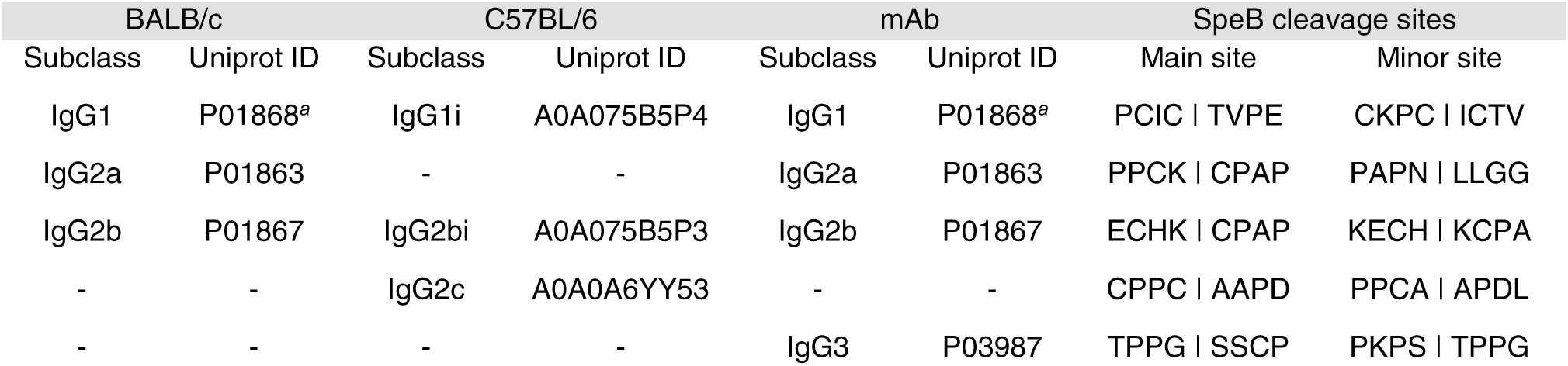
Constant HC regions and SpeB cleavage sites identified in mouse IgGs from different sources. ^*a*^Two amino acid exchanges were detected with respect to the Uniprot entry.

With regard to SpeB induced proteolysis, a main and a minor cleavage site were identified for HC constant regions of each IgG subclass under the applied conditions (Table 1). In accordance with previous reports, the identified cleavage sites are mainly located in unstructured, proline-rich regions, such as the hinge. The resulting C-terminal cleavage products of polyclonal and monoclonal IgGs comprising the glycosylated Fc/2 subunit are listed in Supplementary Table S-1. Based on the identified Fc/2 subunits, we generated extracted ion current chromatograms (XICCs) to assess chromatographic separation of the different cleavage products: indeed, Fc/2 subunits obtained from polyclonal IgGs could be separated in a subclass-specific manner by means of IP-RP-HPLC (Fig. 2a and b). Hence, relative abundances of main and minor C-terminal cleavage products could be determined based on peak areas obtained from the corresponding XICCs (Supplementary Table S-1, Supplementary Fig. S-4).

**Figure 2.**
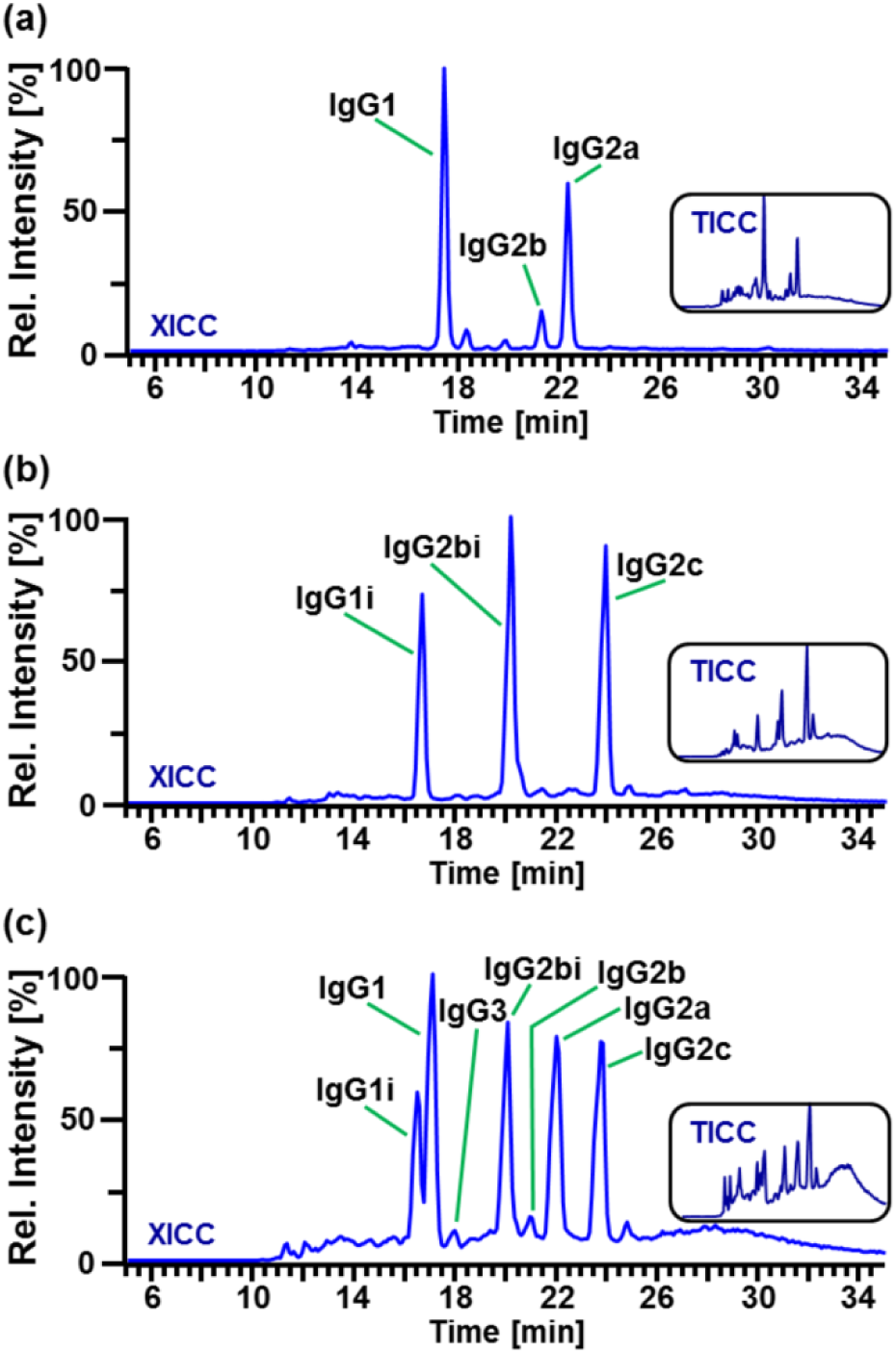
Separation of Fc/2 subunits in polyclonal IgGs from (a) BALB/c and (b) C57BL/6 mice, respectively, and (c) a mixture of polyclonal IgGs from both mouse strains spiked with monoclonal IgG3. Peaks are labelled according to the IgG subclass of the underlying Fc/2 subunit. XICCs represent the sum of signals considering several charge states of two glycoforms (G0F and G1F) for each IgG subclass. Details are provided in the Experimental Section and as Supplementary Excel file. The corresponding TICCs are shown as insets.

In order to further assess the capabilities of the chromatographic method, we analysed a mixture of all described murine IgG subclasses and isotypes, i.e. a combination of polyclonal IgGs from C57BL/6 and BALB/c spiked with monoclonal IgG3. Indeed, Fc/2 subunits from all seven murine IgG subclasses and isotypes could be separated and identified (Fig. 2c). Intriguingly, Fc/2 of even closely related isotypes was readily separated as demonstrated for IgG1 and IgG1i differing in only three out of 213 amino acids. Chromatographic separation is a prerequisite for quantitative analysis of Fc/2 subunits based on XICCs as peak areas may be biased upon co-elution.

### Relative quantification of IgG subclasses

IgG subclasses may be quantified based on relative abundances of the respective Fc/2 subunits. For this purpose, we determined peak areas in corresponding XICCs and corrected relative abundances of Fc/2 for minor cleavage products (Supplementary Table S-1, Supplementary Excel file). The resulting IgG subclass abundance profiles obtained for a BALB/c and a C57BL/6 mouse individual are shown in Fig. 3. Intriguingly, IgG1 predominated in the BALB/c mouse, whereas higher levels of IgG2c (homologous to IgG2a in BALB/c) and IgG2bi were observed in the C57BL/6 individual. This is in accordance with the default immune polarization of the two prototypical mouse strains, arising from T helper type 2 (Th2) and T helper type (Th1) dominated immune responses, respectively.^44^ In contrast to ELISA usually providing comparative values of a specific subclass in multiple samples, the here described IgG subclass profiles enable direct comparison of different subclasses within a mouse individual. This may be favourable for the determination of frequently used IgG1/IgG2a ratios avoiding potential bias arising from differences in the quality of secondary antibodies.^45,46^ Subclass quantification of several mouse individuals was performed in the course of a vaccination study in BALB/c mice (see below).

**Figure 3.**
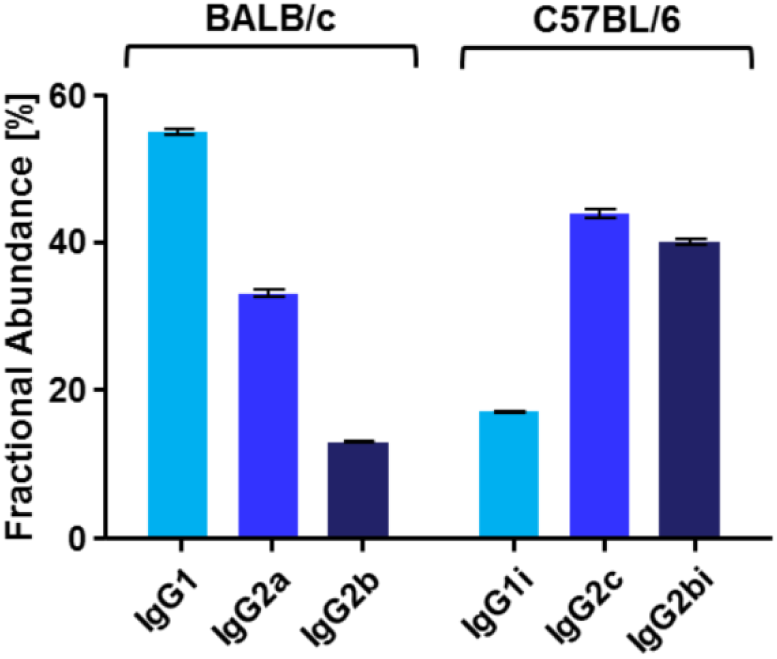
Fractional subclass abundances of total IgGs from BALB/c and C57BL/6 mouse individuals. Relative abundances were obtained from XICCs of Fc/2 subunits including all detected glycoforms (Supplementary Excel file). Mean values are based on three technical replicates; error bars indicate standard deviations.

### Identification and relative quantification of Fc/2 glycoforms

Upon subclass-specific separation and identification of Fc/2 subunits from polyclonal IgGs of BALB/c mice, we assessed *N*-glycosylation variants of Fc/2 based on full scan mass spectra in a middle-up approach. Taking into account the mass of the amino acid sequence as well as the mass of potential *N*-glycan structures, glycoforms were assigned using the MoFi software. Six *N*-glycan structures of the biantennary, complex type were identified for Fc/2 from IgG1 (Fig. 4a and Supplementary Fig. S-5). The assigned structures all comprise core-fucose residues and are substituted with up to two *N*-glycolylneuraminic acid (Neu5Gc) moieties. The same glycans, however at different abundances, were identified for Fc/2 from IgG2a and IgG2b (Supplementary Fig. S-5). In addition, we observed a series of signals shifted by +16 Da with respect to the detected glycoforms. These masses may be attributed to oxidation variants, which is further supported by the detection of oxidised tryptic peptides within Fc/2 (Fig.4a and Supplementary Fig. S-6).

**Figure 4.**
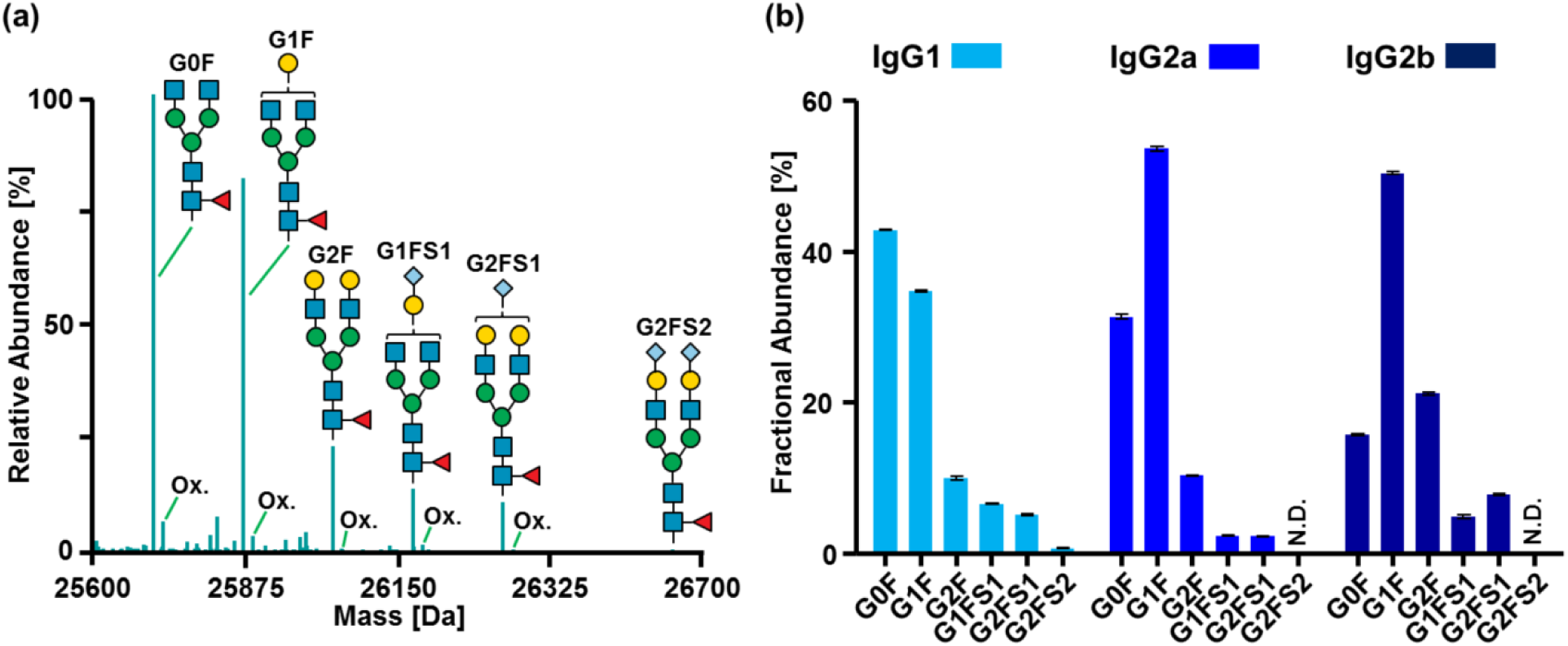
Fc/2 glycoform quantification of polyclonal IgGs from a BALB/c mouse individual. (a) Deconvoluted mass spectrum of Fc/2 from IgG1 with assigned *N*-glycan structures and oxidation variants (Ox.). (b) Fractional abundances of *N*-glycoforms for IgG subclasses. Abbreviations for *N*-glycans correspond to structures shown in (a). Relative abundances were obtained from XICCs of Fc/2 subunits (Supplementary Excel file). Mean values are based on three technical replicates; error bars indicate standard deviations. Raw spectra are shown in Supplementary Fig. S-5.

For relative quantification, peak areas were obtained from XICCs of the identified glycoforms. In IgG1, the non-galactosylated G0F structure predominated whereas the *N*-glycan extended with one galactose residue (G1F) was most abundant in both IgG2a and IgG2b (Fig. 4b). The here described subclass-specific differences of IgG *N*-glycan profiles in naive BALB/c mice are in line with previous reports.^30,40^ In addition to polyclonal IgGs, we analysed monoclonal antibodies of different subclasses applying the above described middle-up approach. Mass spectra revealed additional PTMs in these IgGs: based on characteristic mass shifts, oxidation variants of Fc/2 glycoforms were detected in all analysed mAbs and are exemplified for monoclonal IgG1 and IgG2b in Supplementary Fig. S-7. Moreover, we observed Fc/2 variants in monoclonal IgG1 and IgG2b comprising a mass shift of +128 Da, which may be attributed to incomplete processing of the C-terminal lysine residue. Detection of lysine variants was even possible if main and minor cleavage products differed by one N-terminal lysine residue as demonstrated for IgG2b (Supplementary Fig. S-7). Simultaneous determination of Fc/2 glycosylation and oxidation variants may be of relevance in quality assessment of functional mouse IgGs. Indeed, both modifications impact downstream effector functions, as decreased affinity of oxidised mAbs to Fcγ receptors has been reported.^47^ Thus, profiling of Fc/2 glycosylation and oxidation variants of murine mAbs may be especially relevant for immunological studies in mice using these mAbs as functional tools, e.g. for in vivo depletion of specific cell types via ADCC and complement-dependent cytotoxicity.

### Subclass-specific analysis of Fc/2 from BALB/c mice upon vaccination with Phl p 6

In order to assess the applicability of the described middle-up approach in an immunological context, we set out to determine the effects of a plant allergen on IgG subclass abundances and glycosylation in mice. Specifically, we studied the immunological response of BALB/c mice to vaccination with Phl p 6, a protein allergen from timothy grass pollen (*Phleum pratense*).^33^ Four cohorts of five mouse individuals each were analysed: (i) naive, (ii) Phl p 6 vaccinated, (iii) Phl p 6 co-administered with the Th2 adjuvant aluminumhydroxid (alum), and (iv) Phl p 6 co-administered with the Th1 adjuvant CpG-ODN1826 (CpG). Polyclonal IgGs were obtained from each mouse individual by protein G affinity chromatography. Additionally, antigen-specific IgGs of the three vaccinated mouse groups were affinity-purified from the obtained total IgGs using immobilized Phl p 6. All three fractions, i.e. total, antigen-specific and non-antigen-specific IgGs, were subjected to middle-up analysis for comprehensive characterization of IgG subclass abundances and glycosylation (Supplementary Fig. S-8). Indeed, relative abundances of IgG subclasses drastically changed in response to vaccination with Phl p 6 (Supplementary Fig. S-9). Immunization with Phl p 6 exclusively induced IgG1 antibodies resulting in a significant increase in the relative abundance of total IgG1. Similarly, vaccination with Phl p 6 and the adjuvant alum induced antigen-specific IgG1 and shifted the ratio of total IgG1 to IgG2a/b towards IgG1. In contrast, higher levels of antigen-specific IgG2a and IgG2b were observed upon co-administration of Phl p 6 and the adjuvant CpG. This is in accordance with non-methylated CpG motifs in bacterial or viral DNA promoting Th1 immunity resulting in a class switch to IgG2a.^48^ Intriguingly, relative subclass abundances of non-antigen-specific IgGs differed from those of total IgGs in naive mice only if adjuvants was co-administered with Phl p 6. This effect may be attributed to bystander activation of non-Phl p 6 Th cells accompanied by a class-switch in B cells.^49^

In order to assess the effect of vaccination on *N*-glycosylation profiles, we compared relative glycoform abundances of IgGs from the different mouse cohorts. In case of total IgG1, we observed increased levels of the G1F glycoform and decreased levels of the G0F variant in vaccinated mouse cohorts compared to naive mice, independent of the adjuvants (Supplementary Fig. S-10). In contrast, relative glycoform abundances of total IgG2a and IgG2b were not affected by vaccination as compared to untreated mice. Although absolute changes in glycoform abundances were low, they may be relevant in terms of effector functions. This has previously been described for human IgGs in a HIV vaccination study, where small differences in glycosylation were associated with pronounced changes in antibody effector functions.^26^

To comprehensively compare IgG1 glycosylation patterns of naive mice and the cohort of Phl p 6 treated mice in absence of adjuvant, we generated integrated multivariate glycan profiles using principle component analysis (PCA). Indeed, glycosylation patterns of total IgG1 clustered into groups of immunized and naive mice (Fig. 5a). One mouse of the naive cohort clustered with the immunized group, indicating an ongoing immune reaction, which may occur in specific-pathogen free (SPF) animals. This is supported by elevated levels of total IgG purified from this mouse individual (data not shown). Furthermore, Fc/2 *N*-glycoform abundances of the outlier resemble that of vaccinated mice rather than the naive mouse individuals (Supplementary Fig. S-11).

**Figure 5.**
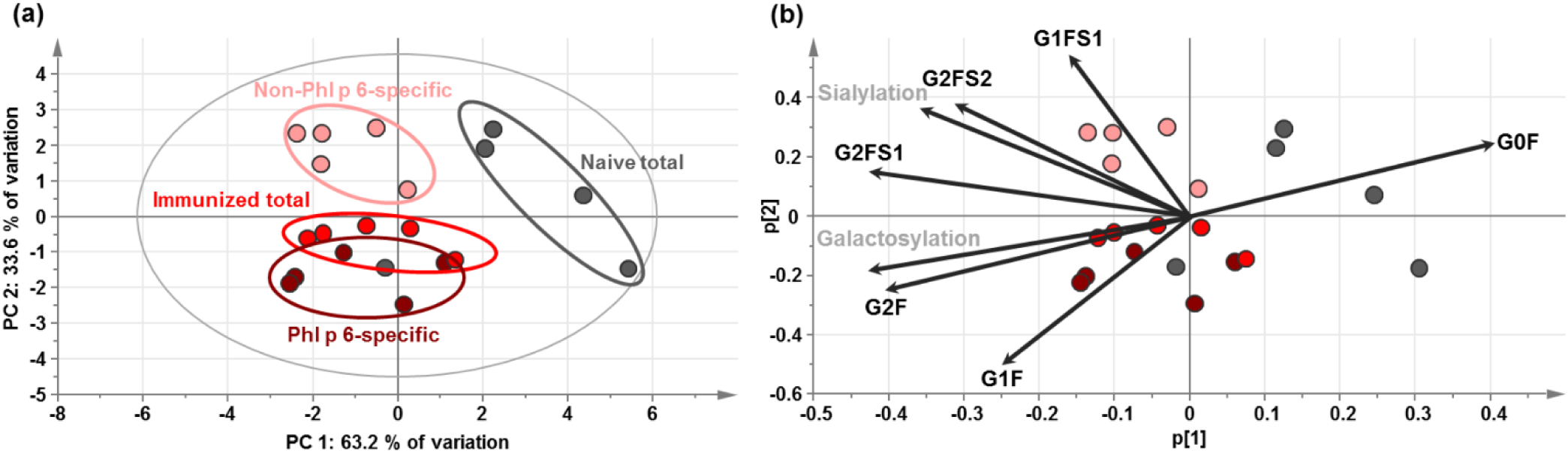
Principal component analysis (PCA) of IgG1 *N*-glycosylation profiles in naive BALB/c mice and BALB/c vaccinated with Phl p 6 in absence of adjuvant. (a) Score plot of IgG1 glycan profiles in total IgG from naive mice (grey) and in three fractions of Phl p 6 vaccinated mice, i.e. total IgG1 (red), Phl p 6-specific IgG1 (dark red) and non-Phl p 6-specific IgG1 (light red). Each dot represents the glycan profile of a mouse individual obtained as mean of three technical replicates. (b) Loading plot showing the contribution of the individual *N*-glycan structures (black) and traits (grey); sialylation and galactosylation refer to the summed weighted abundances of all sialylated and galactosylated structures, respectively, as previously described.^50^ Fractional glycoform abundances are listed in the Supplementary Excel file.

We next compared glycosylation patterns of Phl p 6-specific IgG1 and the remaining fraction of non-Phl p 6-specific IgG1 in immunized mice. Clustering of these two fractions in PCA revealed distinct glycan profiles of antigen-specific and non-antigen-specific IgG1. Intriguingly, all three fractions of the immunized cohort separated from total IgG1 of naive mice in PCA. As illustrated by the corresponding loading plot, the two mouse cohorts mainly differed in their galactosylation levels (Fig. 5b). Separation of Phl p 6-specific and non-Phl p 6-specific IgG1 of the vaccinated cohort can primarily be attributed to differences in the levels of G1F and G1FS1 glycoforms (Fig. 5b). Furthermore, an increase in G1F and G2F structures was observed for Phl p 6-specific IgG1 as compared to non-Phl p 6-specific IgG1. Interestingly, elevated galactosylation levels have been described for antigen-specific IgGs upon immunization with influenza or tetanus vaccine in humans.^50^

## Conclusion

Top-down and middle-up/-down analytical strategies are widely applied for monoclonal antibody characterization. Although highly relevant in immunological studies, analysis of their native counterparts, i.e. polyclonal IgGs, is challenged by immense molecular heterogeneity arising from sequence variability. On the other hand, conservation of the Fc region allows definition of IgG subclasses, which differ in their effector functions and their glycosylation patterns.^51^ Limited sequence variability of Fc may thus enable middle-up analysis of this functionally important IgG subunit. In this study, we explore feasibility of subunit analysis in highly complex mixtures of polyclonal murine IgGs for subclass-specific determination of IgG glycosylation patterns. In order to obtain Fc/2 of all IgG subclasses described in mice, we initially assessed the potential of the bacterial protease SpeB for cleavage in the IgG hinge region. Identification of defined proteolytic products from all IgG subclasses laid the foundation for identification and relative quantification of subclasses and their glycoforms.

The described middle-up approach provides new means to fully assess glycosylation patterns of closely related IgG isotypes, which were not amenable by conventional bottom-up analysis. While tryptic glycopeptides from murine IgGs allow discrimination of subclasses 1, 2 and 3, these specific peptides are isobaric for the four IgG2 isotypes. Although tryptic glycopeptides of IgG2a/2c may be chromatographically separated from those of IgG2b/2bi, discrimination of IgG2a from IgG2c and IgG2b from IgG2bi is not possible because of identical tryptic glycopeptides.^30^ Middle-up analysis of polyclonal IgGs circumvents these restrictions and reveals glycosylation profiles with unlimited subclass specificity. This is especially relevant for immunological studies performed in outbred mouse stocks, e.g. Swiss Webster and CD-1, in which IgG2 isotypes may co-occur.^52^

In addition to revealing subclass-specific glycosylation profiles, we assessed relative subclass abundances. Although a few HPLC-MS based methods have been reported for human IgG subclass quantification at the level of tryptic peptides^53-55^, ELISA is routinely being used to determine specific IgG subclass levels in immunological studies. In this respect, our HPLC-MS based approach overcomes certain limitations of conventional ELISA methods, i.e. requirement for careful standardization when quantitating IgG levels of different subclassed due to differences in the quality of secondary antibodies. Our middle-up approach now provides means to simultaneously quantify IgG1 and IgG2 subclasses comprising six different isotypes in polyclonal murine IgGs thereby providing subclass abundance profiles for each mouse individual. Although readily detectable upon spiking of monoclonal IgG3, Fc/2 subunits of IgG3 were not detected in the investigated polyclonal IgG mixtures, most likely due to the low abundance of this IgG subclass. Downscaling of the applied HPLC-MS setup to the nanoscale may increase sensitivity to enable routine detection of IgG3 and allow the analysis of smaller sample amounts in the future.

Applicability of HPLC-MS based Fc/2 analysis of polyclonal IgGs was demonstrated in a pilot vaccination study of BALB/c mice immunized with the antigen Phl p 6. Antigen-specific IgGs obtained from vaccinated mouse cohorts were found to be predominantly of the IgG1 subclass. With regard to glycosylation, distinct profiles were observed for antigen- and non-antigen-specific IgG1, which were separated even further from total IgG1 of naive mice in multivariate analysis. The observed changes in subclass-specific glycosylation profiles upon vaccination illustrate the potential of global Fc/2 analysis in an immunological context. Comprehensive subclass specific *N*-glycosylation profiling may be applied to study IgG glycosylation in the context of hypersensitive immune responses which is partly governed by IgG binding affinities to different Fcγ receptors.^56^ Furthermore, mouse models for autoimmune diseases, e.g. Sjögren’s syndrome or rheumatoid arthritis, would be compelling topics to be tackled by global Fc/2 profiling.^57,58^

In summary, the described workflow accomplishes global analysis of polyclonal murine IgGs with respect to subclass abundances, glycosylation profiles and occurrence of other modifications, e.g. oxidation and lysine variants. This comprehensive set of information may be obtained in a single analysis involving short sample preparation, standard HPLC-MS analysis, and straightforward data evaluation. Our approach therefore demonstrates the potential of middle-up analysis for polyclonal IgGs and represents an attractive extension to the toolbox of existing bottom-up strategies. In the future, global IgG subclass quantification and *N*-glycoform profiling may be implemented for human polyclonal antibodies at the level of Fc/2 subunits, e.g. for monitoring of IgG subclasses in human vaccination studies.

## Methods

### Materials

Acetonitrile (ACN, ≥ 99.9%) was purchased from VWR International (Vienna, Austria). Ammonium hexa-fluorophosphate (AHFP, 99.99%), trifluoroacetic acid (TFA, ≥ 99.0%), tris(2-carboxyethyl)phosphine (TCEP, ≥ 98.0%), iodoacetamide (IAA, ≥ 99.0%), formic acid (FA, 98.0-100%), L-cysteine HCl (≥ 99.0%), glycine (≥ 99%), sodium chloride (NaCl, ≥ 99.5%), sodium phosphate monobasic (≥ 99.0%), and Gibco® DPBS without calcium and magnesium were obtained from Sigma-Aldrich (Vienna, Austria). Ammonium acetate (≥ 98.0%), ammonia (25% in H2O), acetic acid (≥ 98.5%), ethanol (≥ 99.5%), and sodium hydrogencarbonate (NaHCO3, ≥ 99.5%) were purchased from Merck (Darmstadt, Germany). Tris(hydroxymethyl)amino-methane (TRIS, > 99.9%) and Alu-Gel-S (alum) were obtained from Serva (Heidelberg, Germany). Deionized water was obtained from a MilliQ® Integral 3 instrument (Millipore, Billerica, MA, USA). Ab Spin Trap columns (immobilized protein G) and NHS-activated sepharose® 4 fast flow were purchased from GE Healthcare Life Sciences (Buckinghamshire, UK). Vivaspin® 500 50 kDa MWCO filters were purchased from Sartorius (Göttingen, Germany). Corning® Costar® Spin-X® centrifuge tube filters were obtained from Corning Inc. (Corning, NY, USA). The adjuvant CpG-ODN1826 with phosophothioate backbone (CpG) was purchased from Eurofins Genomics (Ebersberg, Germany). The allergen Phl p 6.0101 (henceforth named Phl p 6) was recombinantly expressed in *Escherichia coli*. Monoclonal murine IgG2a (cat. no. 21380071) and IgG3 (cat. no. 21810201) were purchased from ImmunoTools GmbH (Friesoythe, Germany). SpeB (Streptococcal pyrogenic exotoxin B, FabULOUS®, product number A0-PU1-020) was a kind gift from Genovis AB (Lund, Sweden). Hybridoma supernatant containing monoclonal murine IgG1 and IgG2b were kindly provided by Peter Hammerl (Department of Biosciences, University of Salzburg, Austria). Sequencing grade modified Trypsin was obtained from Promega, Madison, WI, USA.

### Vaccination of BALB/c mice

Female, 6–10 week-old BALB/c and C57BL/6 mice were obtained from Janvier Labs (Le Genest St. Isle, France) and maintained at the animal facility of the University of Salzburg according to local guidelines for animal care. In order to assess changes in Fc-glycosylation upon a vaccination stimulus, four cohorts of BALB/c mice (n=5) were investigated: (i) a naive group, (ii) a Phl p 6 vaccinated group, and two groups vaccinated with Phl p 6 in the presence of the adjuvant (iii) alum or (iv) CpG, respectively. All treated cohorts were vaccinated three times in a two week interval with 10 µg of Phl p 6 in 200 µl of PBS subcutaneously either without adjuvant, or together with 50 µg of CpG or adsorbed to 50% alum. Two weeks after the final vaccination all mice were sacrificed and blood samples were obtained. The blood was allowed to clot for 1.0 h at 21 °C and was subsequently centrifuged to obtain individual serum samples. These samples were sterile-filtered using 0.45 µm filters and stored at - 20 °C until further purification. Animal experiments were approved by the Austrian Ministry of Science (permit number: BMWF-66-012/0024-II/3b/2016).

### Purification of polyclonal IgGs from mouse serum

Serum samples were obtained from BALB/c and C57BL/6 mouse individuals for purification of IgGs. For this purpose, protein G columns were equilibrated five times with 400 µL of 20 mmol.L^-1^ of sodium phosphate buffer (pH 7.0). Four hundred µL of serum were diluted 1:2 with 20 mmol.L^-1^ sodium phosphate buffer (pH 7.0), loaded onto the spin column and incubated for 1.0 h at 21 °C on an overhead rotator. The resin was washed five times with 400 µL of 20 mmol.L^-1^ sodium phosphate buffer (pH 7.0) and bound IgGs were eluted upon incubation with 300 µL of 100 mmol.L^-1^ glycine buffer (pH 2.5) for 15 min at 21 °C on an overhead rotator. The eluate was immediately neutralized by addition of 33 µl of 1.0 mol.L^-1^ TRIS-HCl buffer (pH 8.0). The elution step was repeated twice; eluates were pooled and desalted using a 50 kDa MWCO filter according to manufacturer’s instructions. Purified IgGs were diluted in 175 mmol.L^-1^ ammonium acetate (pH 6.85) to a concentration of 1.0 mg.ml^-1^ for further analysis. IgG concentration was determined at 280 nm using a nanophotometer (model P330, Implen GmbH, München, Germany).

### Purification of Phl p 6-specific antibodies

Antigen-specific IgGs were purified by affinity chromatography using Phl p 6-sepharose beads. For this purpose, Phl p 6 was linked to NHS-activated sepharose according to manufacturer’s instructions. Coupling was performed in 2 mmol.L^-1^ sodium phosphate buffer, pH 7.4 at 4.0 °C on an overhead rotator for 19 h. Residual NHS-groups were blocked with 100 mmol.L^-1^ TRIS-HCl buffer (pH 8.5) for 4.0 hours at 4.0 °C while rotating. Phl p 6-sepharose beads were stored in 20% (v/v) ethanol solution at 4 °C until use. For purification of antigen-specific IgGs, a column was prepared by equilibrating 250 µL of Phl p 6 resin in spin filters five times with 400 µL of PBS (pH 7.4). Protein G-purified total IgGs were diluted 1:2 (v/v) with PBS (pH 7.4), mixed with the equilibrated beads and incubated for 2 h at 21 °C on an overhead rotator. After washing the resin five times with 400 µL of PBS (pH 7.4), antigen-specific IgGs were eluted, buffer exchanged and diluted as described above.

### Sample preparation

For SpeB proteolysis, monoclonal or polyclonal IgGs were diluted in 175 mmol.L^-1^ ammonium acetate to a concentration of 0.3 mg.mL^-1^ and supplemented with 50 mmol.L^-1^ L-cysteine. Digestion with SpeB was performed at an enzyme to substrate ratio of 2 U.µg^-1^ for 3.0 h at 37 °C while shaking at 850 rpm. Samples were subjected to HPLC-MS analysis without further purification. For analysis of tryptic peptides, protein G-purified polyclonal IgGs from BALB/c were reduced with 5 mmol.L^-1^ TCEP for 15 min at 60 °C in 175 mmol.L^-1^ ammonium acetate at 0.1 mg.mL^-1^ and alkylated with 20 mmol.L^-1^ IAA for 30 min at 22 °C in the dark. Tryptic digestion was performed at a substrate to enzyme ratio of 50:1 (*w/w*) for 3.0 h at 37 °C.

### High-performance liquid chromatography

Chromatographic separation was carried out on a capillary HPLC instrument (UltiMate™ U3000 RSLC, Thermo Fisher Scientific, Germering, Germany). SpeB digests were separated on an AdvanceBio Diphenyl column featuring superficially porous particles (150 × 2.1 mm i.d., 3.5 μm particle size, 450 Å pore size, Agilent, Santa Clara, CA, USA) at a flow rate of 200 µL.min^-1^ and a column oven temperature of 80 °C. Ten microliters of sample [0.3 mg.mL^-1^] were injected using in-line split-loop mode. A linear gradient of mobile phase solutions A (H2O + 0.050% TFA) and B (ACN + 0.050% TFA) was applied as follows: 20.0% B for 5.0 min, 27.5%–40.0% B in 30 min, 80.0% B for 5.0 min, and 20.0% B for 20 min. UV-detection was carried out at 214 nm using a 1.4 µl flow cell. Tryptic peptides were analysed using a Hypersil GOLD aQ C18 column (100 × 1.0 mm i.d., 1.9 µm particle size, 175 Å pore size, Thermo Fisher Scientific, Sunnyvale, CA, USA) at a flow rate of 100 μL.min^−1^ and a column oven temperature of 50 °C applying the following gradient: mobile phase A (H2O + 0.10% FA), mobile phase B (acetonitrile + 0.10% FA); 2.0% B for 5 min, 2.0−12.0% B in 5 min, 12.0−45.0% B in 40 min, 80% B for 5 min, and 2.0% B for 25 min. Ten microliters of sample [0.1 mg.mL^-1^] were injected using in-line split-loop mode.

### Mass spectrometry

SpeB digests were analysed on a Thermo Scientific™ QExactive™ benchtop quadrupole-Orbitrap™ mass spectrometer equipped with an Ion Max™ source with a heated electrospray ionization (HESI) probe, both from Thermo Fisher Scientific (Bremen, Germany), and an MXT715-000 - MX Series II Switching Valve (IDEX Health & Science LLC, Oak Harbor, WA, USA). Mass spectrometric data were acquired similar to a previous study.^10^ Specifically, the following instrument settings were used: source heater temperature of 250 °C, spray voltage of 3.5 kV, sheath gas flow of 30 arbitrary units, auxiliary gas flow of 10 arbitrary units, capillary temperature of 320 °C, in-source collision-induced dissociation (IS-CID) of 0.0 eV, S-lens RF level of 80.0, AGC target of 3e6 at *m/z* of 1500–3000 with a maximum injection time of 200 ms, resolution of 140,000 at 200 *m/z*, and averaging of 10 microscans. MS/MS was carried out in three runs applying all ion fragmentation (AIF) in the higher-energy collisional dissociation (HCD) cell at normalized collision energy (NCE) settings of 18.0, 20.0, and 22.0, respectively, within a scan range of *m/z* 500–3,000 (corresponding to 63, 70, and 77 eV collision energy for a singly charged ion at *m/z* 2,000), and a resolution setting of 140,000 at *m/z* 200. MS analysis of tryptic peptides was performed on a Thermo Scientific™ Q Exactive™ Plus benchtop quadrupole-Orbitrap™ mass spectrometer equipped with an Ion Max™ source with a HESI probe, both from Thermo Fisher Scientific (Bremen, Germany). Instrument settings were as follows: source heater temperature of 100 °C, spray voltage of 3.5 kV, sheath gas flow of 10 arbitrary units, auxiliary gas flow of 5 arbitrary units, capillary temperature of 300 °C, S-lens RF level of 60.0. Each scan cycle consisted of a full scan at a scan range of *m/z* 350-2,000 with an AGC target of 1e6, a maximum injection time of 150 ms and a resolution setting of 70,000, followed by 5 data-dependent HCD scans at 28 NCE (corresponding to 27.5 eV for the selected peptide) with an AGC target of 5e5, a maximum injection time of 150 ms and a resolution setting of 17,500. Both instruments were mass calibrated using Pierce™ LTQ Velos ESI Positive Ion Calibration Solution from Life Technologies (Vienna, Austria) and AHFP.

### Data evaluation

Isotopically resolved mass spectra were deconvoluted using the Xtract algorithm implemented in the Xcalibur™ software version 3.0.63 (Thermo Fisher Scientific, Waltham, MA, USA). Annotation of glycoforms and other PTMs was conducted using MoFi v1.0.^59^ Fc/2 amino acid sequences were derived from constant heavy chain entries in the Uniprot database (Supplementary Fig. S-2). MoFi-assisted peak assignment was based on monoisotopic masses with a mass tolerance of ±50 ppm assuming intrachain disulphide bridges to be intact and interchain disulphide bonds to be reduced (see amino acid sequences Supplementary Fig. S-2). A library containing the six most common *N*-glycan structures reported for murine IgGs and their afucosylated counterparts was used (Supplementary Table S-2). For relative quantification of Fc/2 *N*-glycoforms, extracted ion current chromatograms (XICCs) were generated and evaluated using the Chromeleon™ Chromatography Data System, version 7.2 SR3 (Thermo Fisher Scientific, Waltham, MA, USA), as previously described^10^: XICC ranges were determined based on a simulated isotopic distribution for the atomic composition of Fc/2 plus the respective glycan taking into account the ten most abundant isotope peaks. Spectra were simulated using the Xcalibur™ software. XICCs were then generated for charge states 10+ to 15+ for relative quantification of subclasses and Fc/2 glycoforms based on integrated peak areas of these XICCs. Fc/2 masses were calculated using the software tool GPMAW 9.51 rel. 0314 (Lighthouse Data, Odense, Denmark).^60^ AIF data were evaluated using the software ProSight Lite v1.4 Build 1.4.6 provided by the Kelleher Research Group (Northwestern University, Evanston, IL, U.S.A.)^61^; mass tolerance for annotation of b- and y-fragments was set to 25 ppm. For statistical evaluation and plotting of data we used GraphPad Prism^®^ version 5.01. Principal component analysis was performed with Simca® 13.0.3 (Umetrics, Sartorius Stedim Biotech, Göttingen, Germany) using unit variances and mean centering for data pre-processing.

### Data availability

The mass spectrometry proteomics data have been deposited to the ProteomeXchange Consortium via the PRIDE partner repository with the dataset identifier PXD014710.^62^

## Acknowledgements

The financial support by the Austrian Federal Ministry of Science, Research, and Economy and by a Start-up Grant of the State of Salzburg is gratefully acknowledged. C. B. acknowledges a research stipend form the István and Agnes Halász Foundation, Saarbrücken, Germany. This work was supported by a grant from the Austrian Science Fund (FWF, W1213). We thank Peter Hammerl and Angelika Stöcklinger (Department of Biosciences, University of Salzburg) for animal handling and scientific discussions and Wolfgang Esser-Skala (Department of Biosciences, University of Salzburg) for scientific discussions. We are grateful to Genovis AB (Lund, Sweden) for providing the SpeB protease. The Open Access Publication Fund of the University of Salzburg is acknowledged for financial support.

## Author contributions statement

C.B., C.R., C.G.H., R.W., and T.W. conceived the study. C.B., C.R., and P.W. performed the experiments; C.B., R.W., and T.W. analysed and interpreted the data. C.B., C.G.H., and T.W. drafted the manuscript. All authors reviewed the manuscript.

## Additional information

The authors declare the following competing financial interests: The salary of Therese Wohlschlager is fully funded; Christian G. Huber’s salary is partly funded by the Christian Doppler Laboratory for Biosimilar Characterization which receives financial support from Novartis and Thermo Fisher Scientific. The authors declare no other conflict of interest.

