## Supplementary Information for "Towards middle-up analysis of polyclonal antibodies: subclass-specific *N*-glycosylation profiling of murine immunoglobulin G (IgG) by means of HPLC-MS"

### Table of contents

Figure S-1: Schematic workflow for SpeB cleavage site determination in HC portions — Page S-2

Figure S-2: Amino acid sequences of HC constant regions of murine IgG subclasses — Page S-3

Figure S-3: Fragment ion spectrum of tryptic peptide [269-286] of IgG1 from BALB/c — Page S-4

Figure S-4: Separation and quantification of main and minor Fc/2 cleavage products — Page S-5

Figure S-5: Raw mass spectra of IgG1, IgG2a and IgG2b from BALB/c — Page S-6

Figure S-6: Identification of oxidised tryptic peptide [179-194] of IgG1 from BALB/c — Page S-7

Figure S-7: Deconvoluted mass spectra of Fc/2 from monoclonal IgG1 and IgG2b — Page S-8

Figure S-8: Schematic workflow for Phl p 6 vaccination study — Page S-9

Figure S-9: IgG subclass abundances after Phl p 6 vaccination — Page S-10

Figure S-10: Subclass-specific glycosylation profiles of total IgG after Phl p 6 vaccination — Page S-11

Figure S-11: Glycoform abundances of IgG1 from mouse individuals — Page S-12

Table S-1: SpeB cleavage products of murine IgG subclasses — Page S-13

Table S-2: *N*-glycan structures considered in glycoform assignment — Page S-14

SI References — Page S-15

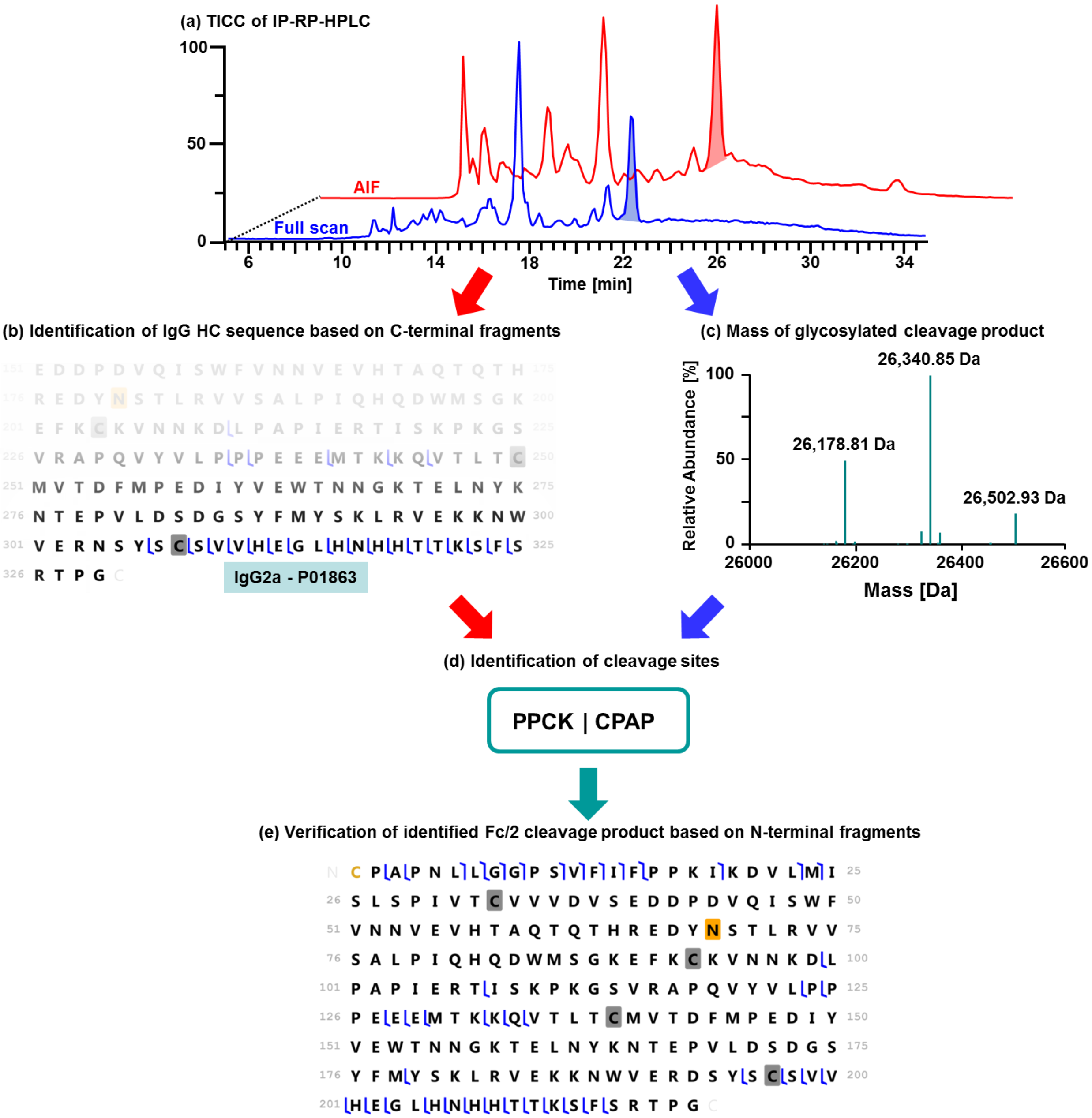


**Figure S-1.** Schematic workflow for SpeB cleavage site determination in HC portions of IgG as shown for polyclonal IgG2a (P01863) from BALB/c. (a) SpeB cleavage products are separated by IP-RP-HPLC and detected in full MS/AIF mode. The AIF trace is depicted at a time offset of 4.0 min. (b) Fragment spectra obtained by AIF are deconvoluted and searched against the sequences provided in Fig. S-2 using the ProSight software.[1] The constant HC sequence is identified based on the assigned y-ions, i.e. C-terminal fragments (blue marks). (c) Masses of the glycosylated cleavage product are derived from the corresponding full scan spectra. (d) The cleavage site is determined based on the mass of the obtained proteolytic product taking into account the G1F glycoform. The process is supported by the GPMAW software.[2] (e) For further confirmation, the sequence of the identified proteolytic product is searched against AIF spectra to match b-ions, i.e. N-terminal fragments, using the ProSight software. Fragmentation data for all identified cleavage sites are available as Supplementary Excel file.

**IgG1 (P01868)**

```
1    AKTTPPSVYP LAPGSAAQTN SMVTLGCLVK GYFPEPVTVT WNSGSLSSGV HTFPAVLQSD
61   LYTLSSSVTV PSSPRPSETV TCNVAHPASS TKVDKKIVPR DCGCKPCICT VPEVSSVFIF
121  PPKPKDVLTI TLTPKVTCVV VDISKDDPEV QFSWFVDDVE VHTAQTQPRE EQFNSTFRSV
181  SELPIMHQDW LNGKEFKCRV NSAAFPAPIE KTISKTKGRP KAPQVYTIPP PKEQMAKDKV
241  SLTCMITDFF PEDITVEWQW NGQPAENYKN TQPIMDTDGS YFVYSKLNVQ KSNWEAGNTF
301  TCSVLHEGLH NHHTEKSLSH SPGK
```

**IgG1i (A0A075B5P4)**

```
1    XKTTPPSVYP LAPGSAAQTN SMVTLGCLVK GYFPEPVTVT WNSGSLSSGV HTFPAVLQSD
61   LYTLSSSVTV PSSTWPSQTV TCNVAHPASS TKVDKKIVPR DCGCKPCICT VPEVSSVFIF
121  PPKPKDVLTI TLTPKVTCVV VDISKDDPEV QFSWFVDDVE VHTAQTKPRE EQINSTFRSV
181  SELPIMHQDW LNGKEFKCRV NSAAFPAPIE KTISKTKGRP KAPQVYTIPP PKEQMAKDKV
241  SLTCMITNFF PEDITVEWQW NGQPAENYKN TQPIMDTDGS YFVYSKLNVQ KSNWEAGNTF
301  TCSVLHEGLH NHHTEKSLSH SPGK
```

**IgG2a (P01863)**

```
1    AKTTAPSVYP LAPVCGDTTG SSVTLGCLVK GYFPEPVTLT WNSGSLSSGV HTFPAVLQSD
61   LYTLSSSVTV TSSTWPSQSI TCNVAHPASS TKVDKKIEPR GPTIKPCPPC KCPAPNLLGG
121  PSVFIFPPKI KDVLMISLSP IVTCVVVDVS EDDPDVQISW FVNNVEVHTA QTQTHREDYN
181  STLRVVSALP IQHQDWMSGK EFKCKVNNKD LPAPIERTIS KPKGSVRAPQ VYVLPPPEEE
241  MTKKQVTLTC MVTDFMPEDI YVEWTNNGKT ELNYKNTEPV LDSDGSYFMY SKLRVEKKNW
301  VERNSYSCSV VHEGLHNHHT TKSFSRTPGK
```

**IgG2b (P01867)**

```
1    KTTPPSVYPL APGCGDTTGS SVTLGCLVKG YFPESVTVTW NSGSLSSSVH TFPALLQSGL
61   YTMSSSVTVP SSTWPSQTVT CSVAHPASST TVDKKLEPSG PISTINPCPP CKECHKCPAP
121  NLEGGPSVFI FPPNIKDVLM ISLTPKVTCV VVDVSEDDPD VQISWFVNNV EVHTAQTQTH
181  REDYNSTIRV VSTLPIQHQD WMSGKEFKCK VNNKDLPSPI ERTISKIKGL VRAPQVYILP
241  PPAEQLSRKD VSLTCLVVGF NPGDISVEWT SNGHTEENYK DTAPVLDSDG SYFIYSKLNM
301  KTSKWEKTDS FSCNVRHEGL KNYYLKKTIS RSPGK
```

**IgG2bi (A0A075B5P3)**

```
1    XKTTPPSVYP LAPGCGDTTG SSVTLGCLVK GYFPESVTVT WNSGSLSSSV HTFPALLQSG
61   LYTMSSSVTV PSSTWPSQTV TCSVAHPASS TTVDKKLEPS GPISTINPCP PCKECHKCPA
121  PNLEGGPSVF IFPPNIKDVL MISLTPKVTC VVVDVSEDDP DVRISWFVNN VEVHTAQTQT
181  HREDYNSTIR VVSALPIQHQ DWMSGKEFKC KVNNKDLPSP IERTISKIKG LVRAPQVYIL
241  PPPAEQLSRK DVSLTCLVVG FNPGDISVEW TSNGHTEENY KDTAPVLDSD GSYFIYSKLD
301  IKTSKWEKTD SFSCNVRHEG LKNYYLKKTI SRSPGK
```

**IgG2c (A0A0A6YY53)**

```
1    XKTTAPSVYP LAPVCGGTTG SSVTLGCLVK GYFPEPVTLT WNSGSLSSGV HTFPALLQSG
61   LYTLSSSVTV TSNTWPSQTI TCNVAHPASS TKVDKKIEPR VPITQNPCPP LKECPPCAAP
121  DLLGGPSVFI FPPKIKDVLM ISLSPMVTCV VVDVSEDDPD VQISWFVNNV EVHTAQTQTH
181  REDYNSTLRV VSALPIQHQD WMSGKEFKCK VNNRALPSPI EKTISKPRGP VRAPQVYVLP
241  PPAEEMTKKE FSLTCMITGF LPAEIAVDWT SNGRTEQNYK NTATVLDSDG SYFMYSKLRV
301  QKSTWERGSL FACSVVHEGL HNHLTTKTIS RSLGK
```

**IgG3 (P03987)**

```
1    TTTAPSVYPL VPGCSDTSGS SVTLGCLVKG YFPEPVTVKW NYGALSSGVR TVSSVLQSGF
61   YSLSSLVTVP SSTWPSQTVI CNVAHPASKT ELIKRIEPRI PKPSTPPGSS CPPGNILGGP
121  SVFIFPPKPK DALMISLTPK VTCVVVDVSE DDPDVHVSWF VDNKEVHTAW TQPREAQYNS
181  TFRVVSALPI QHQDWMRGKE FKCKVNNKAL PAPIERTISK PKGRAQTPQV YTIPPPREQM
241  SKKKVSLTCL VTNFFSEAIS VEWERNGELE QDYKNTPPIL DSDGTYFLYS KLTVDTDSWL
301  QGEIFTCSVV HEALHNHHTQ KNLSRSPGK
```

**Figure S-2.** Amino acid sequences of HC constant regions of murine IgG subclasses. Uniprot accession numbers of the secreted isoforms are given in brackets. The main cleavage products obtained upon digestion with SpeB are highlighted in grey; minor cleavage sites are marked with a blue line. *N*-glycosylation sites are highlighted in yellow; the C-terminal lysine residue (underlined) is usually absent in polyclonal IgGs but may be present in mAbs. Cysteine residues of intact intramolecular disulphide bonds identified after SpeB cleavage under reducing conditions are highlighted in light blue. Amino acid exchanges (N to D) identified in IgG1 (P01868) of BALB/c mice are highlighted in green (see fragment spectrum in Fig. S-3).

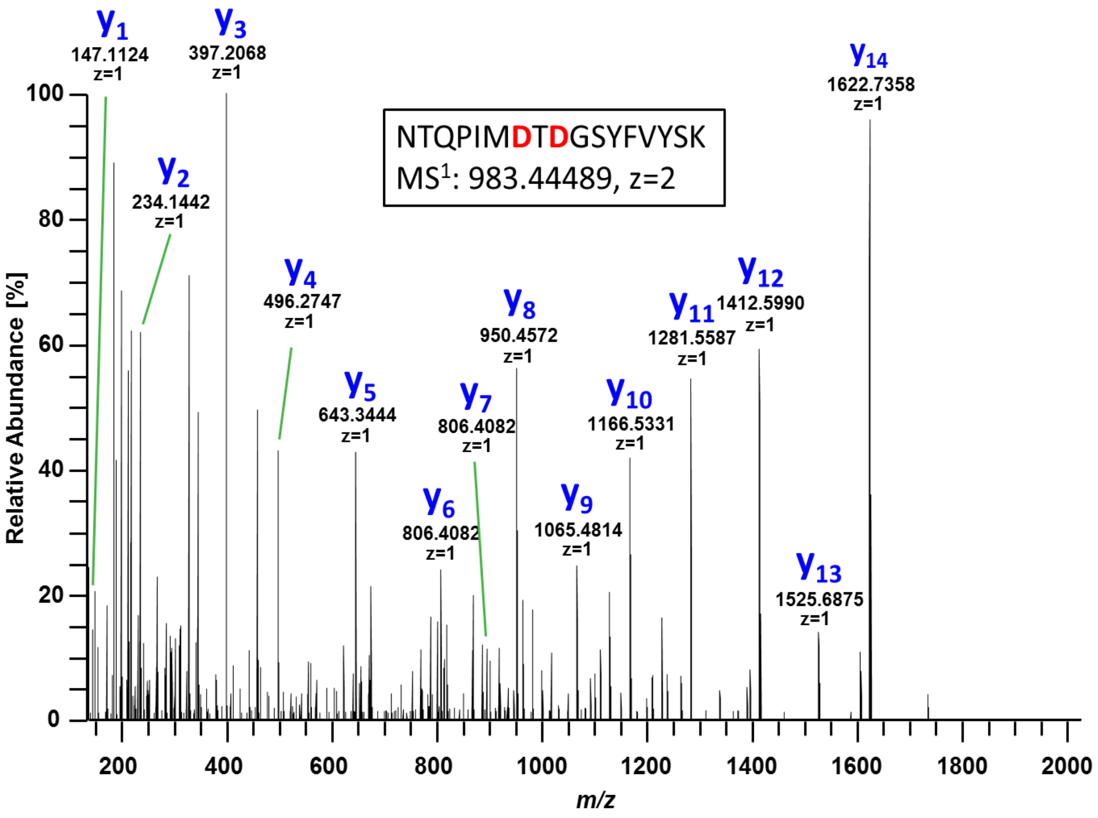


**Figure S-3.** Fragment ion spectrum of tryptic peptide [269-286] from the constant HC region of IgG1 (P01868, Fig. S-2) from BALB/c. Fragmentation was performed at 27.5 eV in the HCD cell; fragment ions were annotated with the help of GPMAW. Amino acid exchanges (red) are in accordance with a sequence conflict of P01868 listed in the Uniprot database.

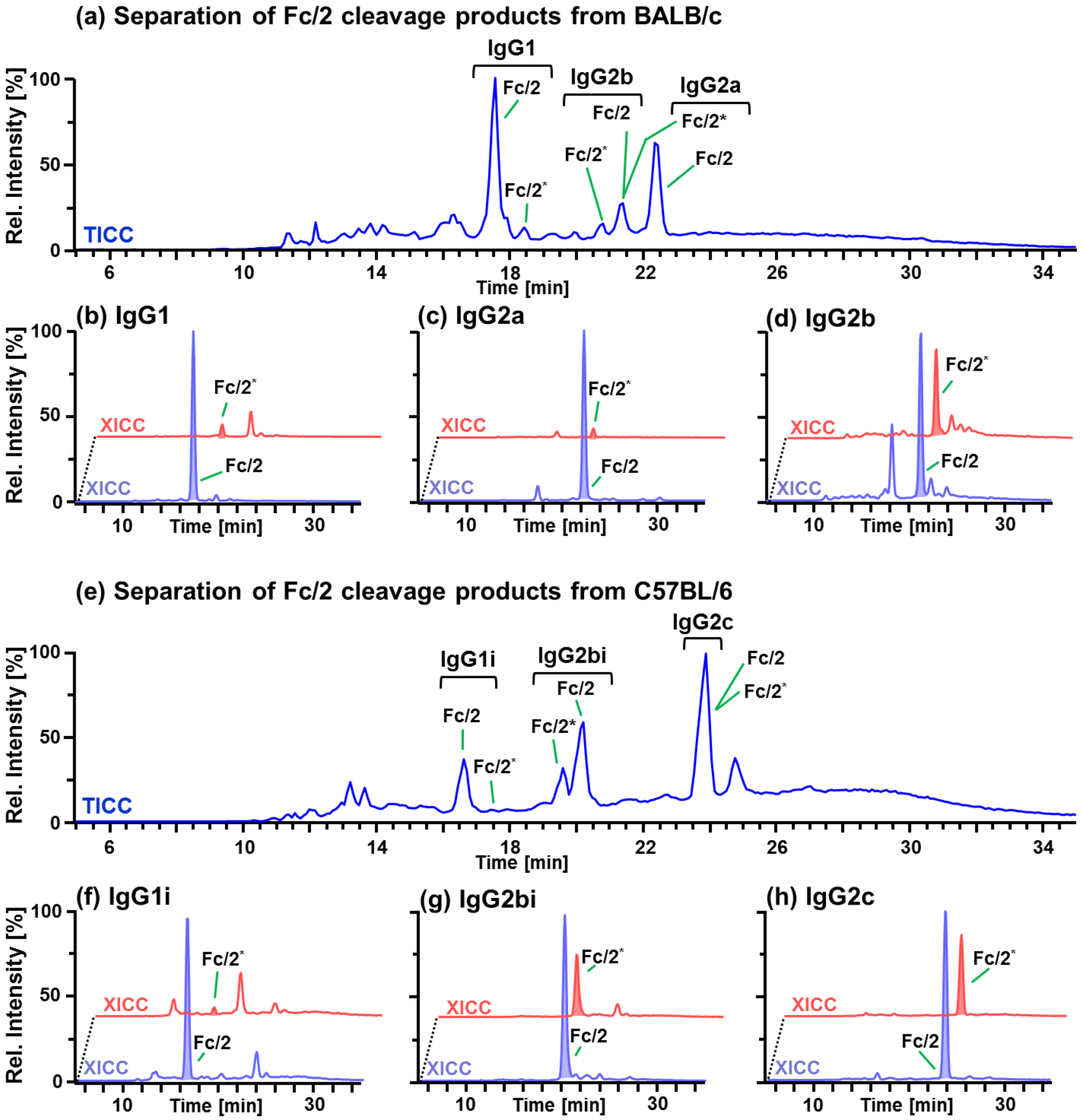


**Figure S-4.** Separation and quantification of Fc/2 cleavage products generated by SpeB. Main and minor cleavage products are designated Fc/2 and Fc/2*, respectively. (a) TICC of SpeB cleavage products of polyclonal IgGs from BALB/c. XICCs generated for Fc/2 and Fc/2* of (b) IgG1, (c) IgG2a and (d) IgG2b. (e) TICC of SpeB cleavage products of polyclonal IgGs from C57BL/6. XICCs generated for Fc/2 and Fc/2* of (f) IgG1i, (g) IgG2bi and (h) IgG2c. All XICCs were generated for G1F *N*-glycoforms lacking the C-terminal lysine residue; intramolecular disulphide bonds were considered to be intact. Peak areas used for relative quantification are indicated. Relative abundances of main and minor cleavage products are listed in Table S-2. XICCs for Fc/2* are depicted at a time offset of 2.0 min. Note that Fc/2 and Fc/2* cleavage products of each IgG subclass are chromatographically separated with the exception of IgG2c.

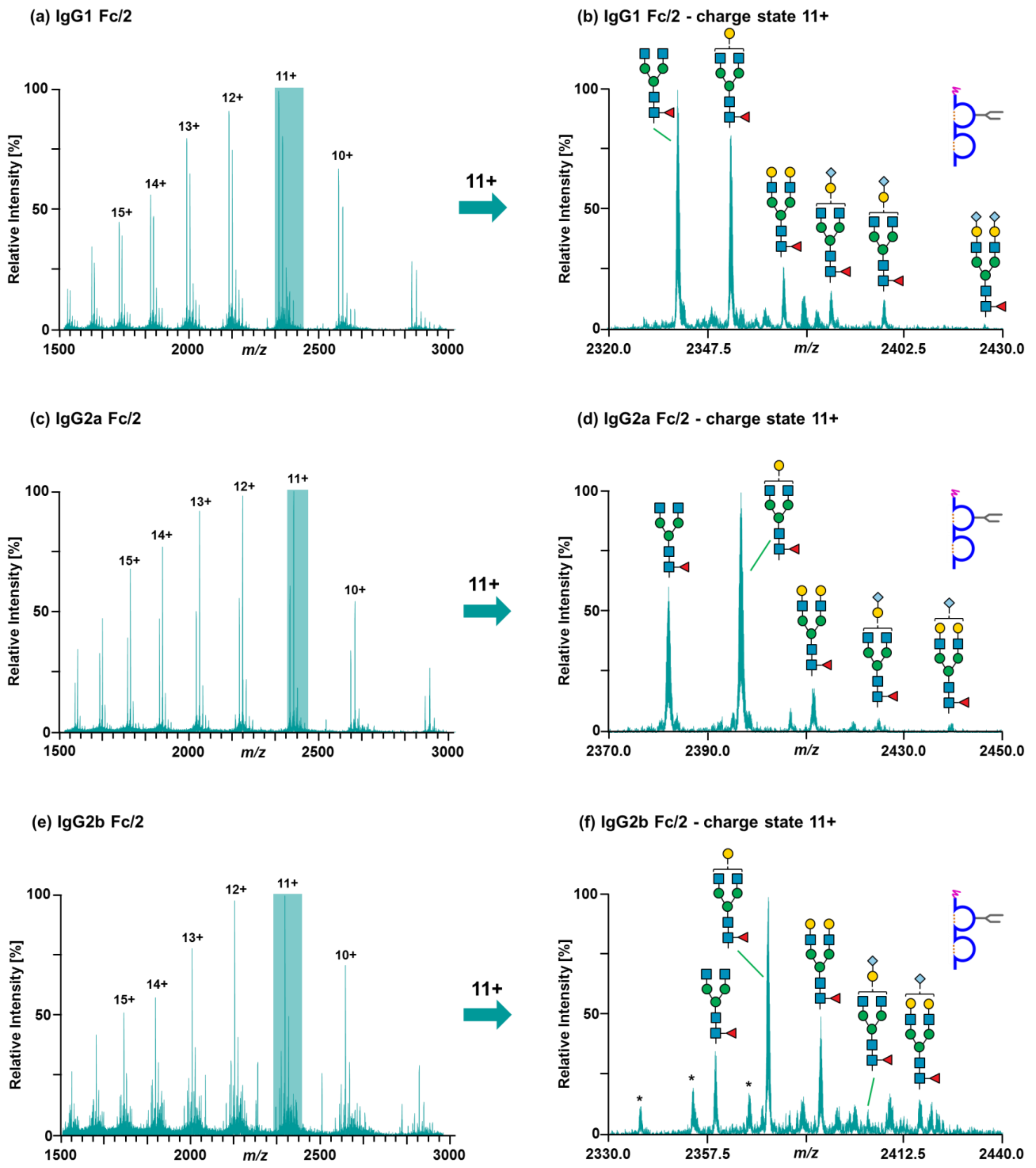


**Figure S-5.** Exemplary raw mass spectra of Fc/2 from (a) IgG1, (c) IgG2a and (e) IgG2b acquired in a total IgG preparation from a naive BALB/c mouse individual. Charge states used for generation of XICCs are indicated. Zooms of the most abundant charge states with assigned *N*-glycoforms are shown (b, d, f). Asterisks in (f) indicate masses assigned to co-eluting Fc/2* from IgG2a (Fig. S-4a).

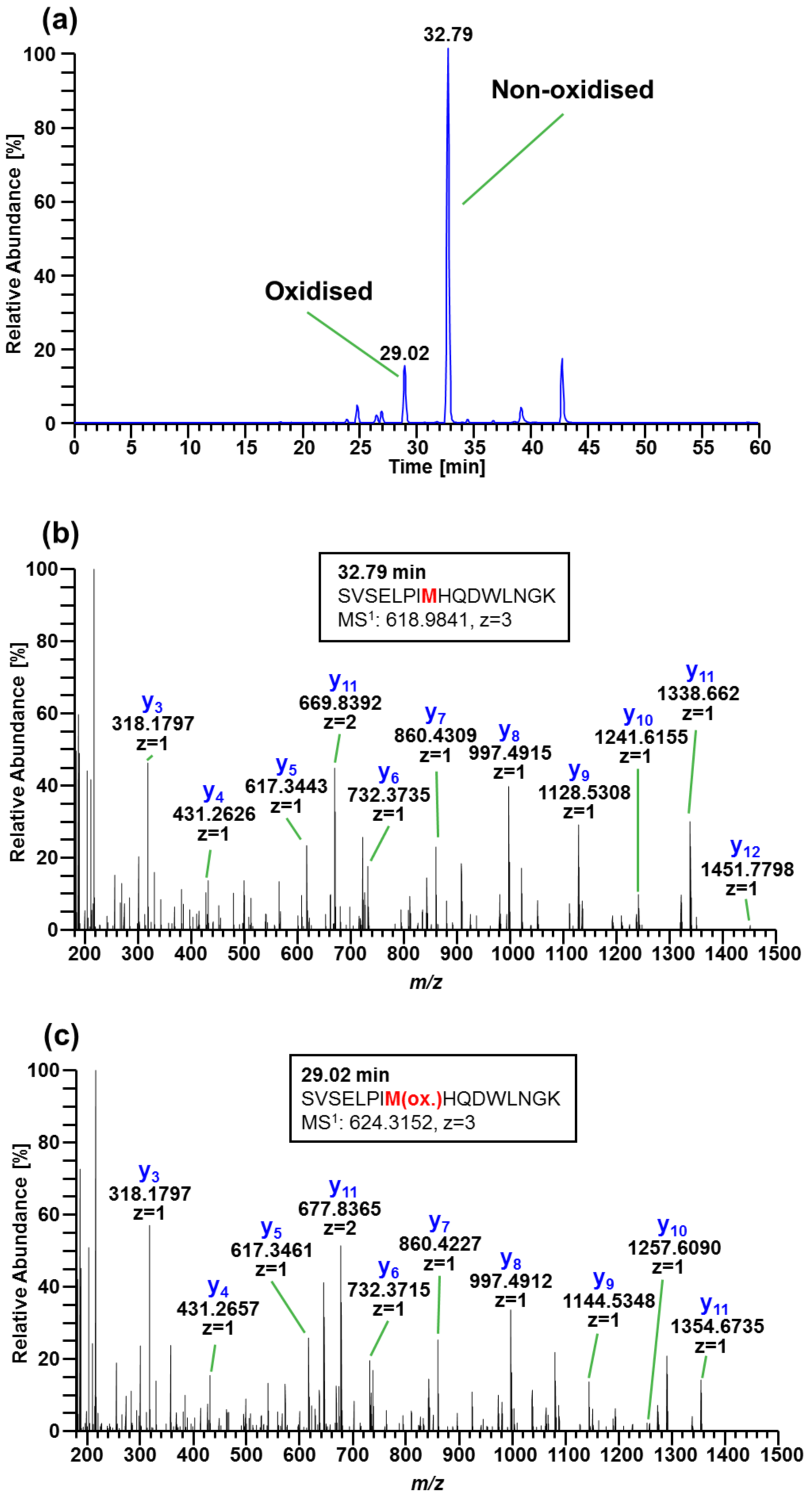


**Figure S-6.** Identification of oxidised tryptic peptide [179-194] from the constant HC region of IgG1 (P01868, Fig. S-2) from BALB/c. (a) XICC of non-oxidised and oxidised version of tryptic peptide [179-194]. The XICC was generated using the Xcalibur™ software taking into account doubly and triply charged ion species (non-oxidised: 618.6453 *m/z* and 927.4644 *m/z*; oxidised: 623.9770 *m/z* and 935.4618 *m/z;* isolation window ± 0.5 *m/z*). (b) Annotated fragment ion spectrum of the non-oxidised variant. (c) Annotated fragment ion spectrum of the oxidised variant. Fragmentation was performed at 27.5 eV in the HCD cell; fragment ions were annotated with the help of GPMAW.

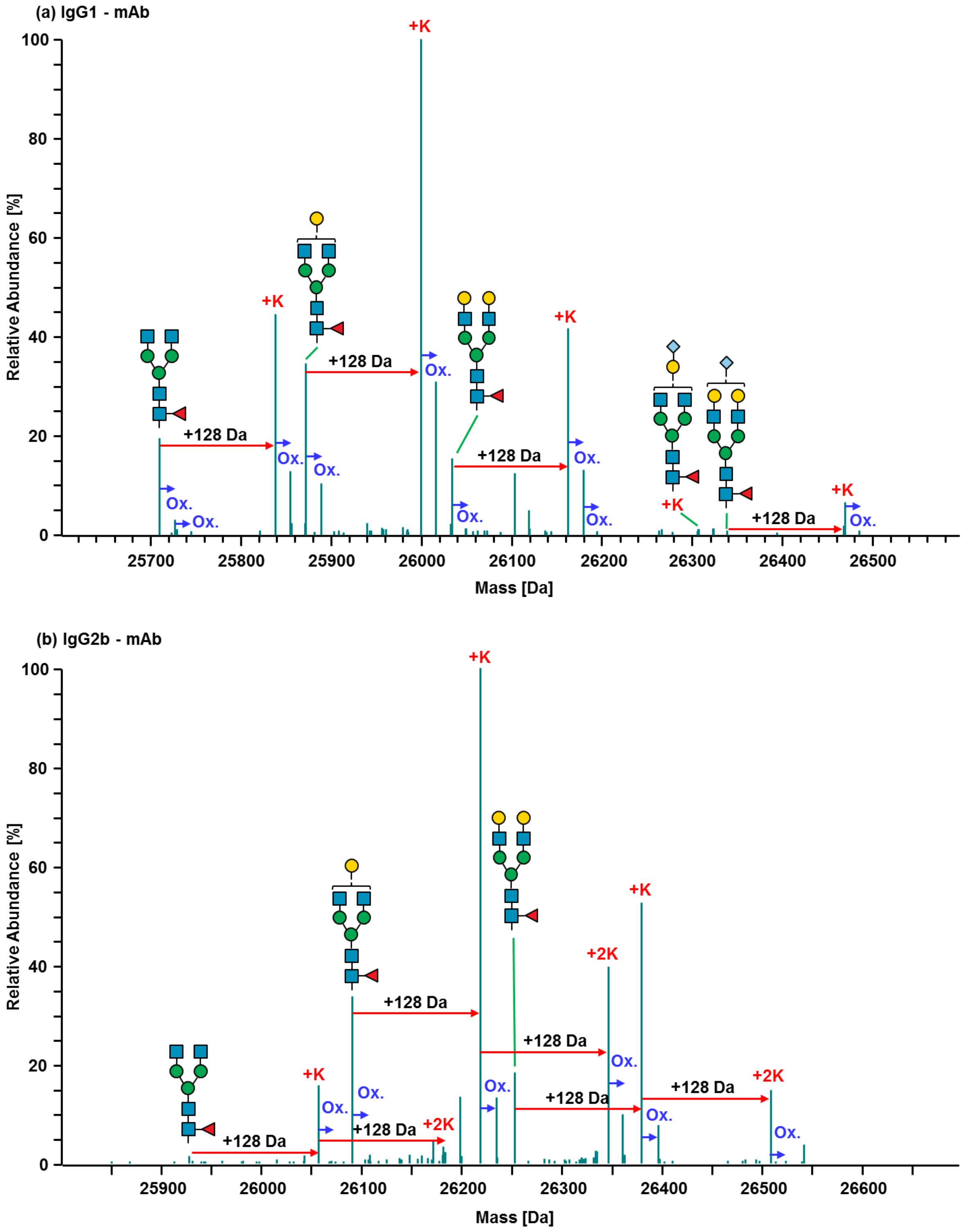


**Figure S-7.** Deconvoluted mass spectra of Fc/2 derived from (a) monoclonal murine IgG1 (P01868) and (b) monoclonal murine IgG2b (P01867). In addition to the main *N*-glycoforms, variants arising from oxidation (Ox.) or comprising additional lysine residues (+K, +2K) were identified based on characteristic mass shifts (+16 Da and +128 Da, respectively).

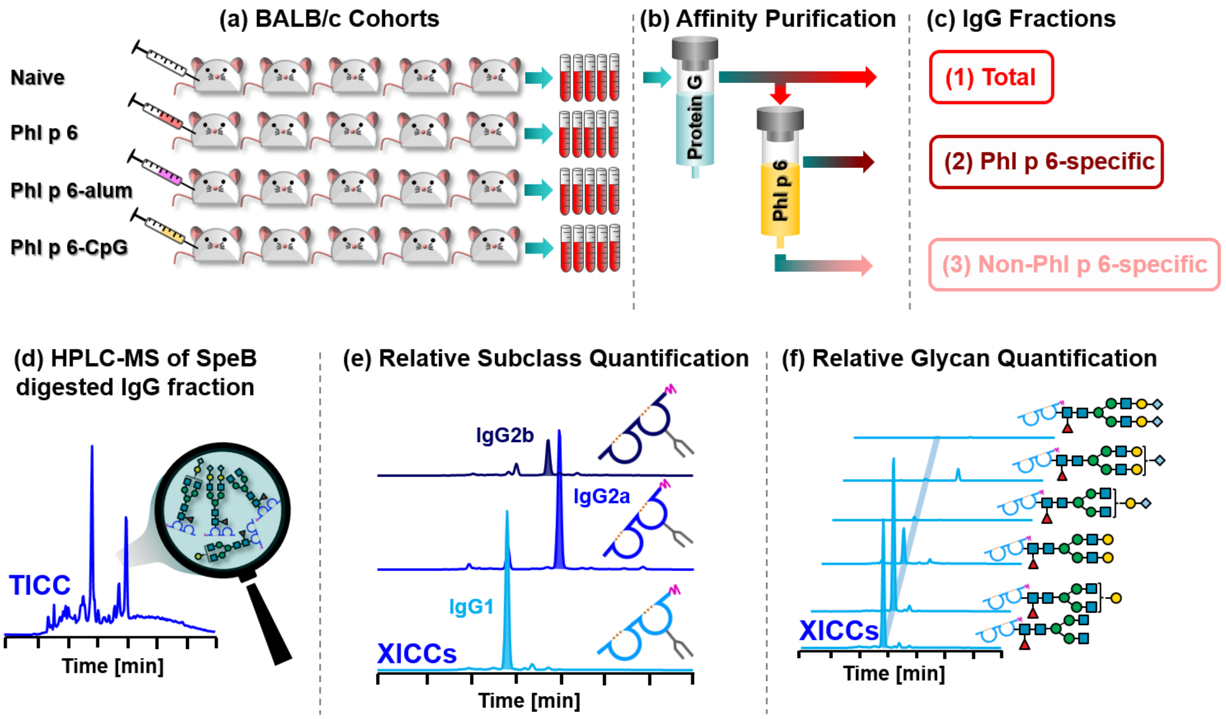


**Figure S-8.** Schematic workflow of the Phl p 6 vaccination study in BALB/c mice. (a) Serum was obtained from naive mice and mice treated with Phl p 6 in the absence or presence of adjuvants (alum or CpG), respectively (n=5). (b) Polyclonal IgGs of all subclasses were purified by protein G affinity chromatography; antigen-specific IgGs of vaccinated mice were affinity-purified from total IgGs using Phl p 6-sepharose. (c) Three IgG fractions were obtained for each vaccinated mouse individual. (d) SpeB digests of IgG fractions were analysed by means of HPLC-MS. XICCs of glycosylated Fc/2 cleavage products were generated to determine (e) relative subclass abundances and (f) relative *N*-glycoform abundances.

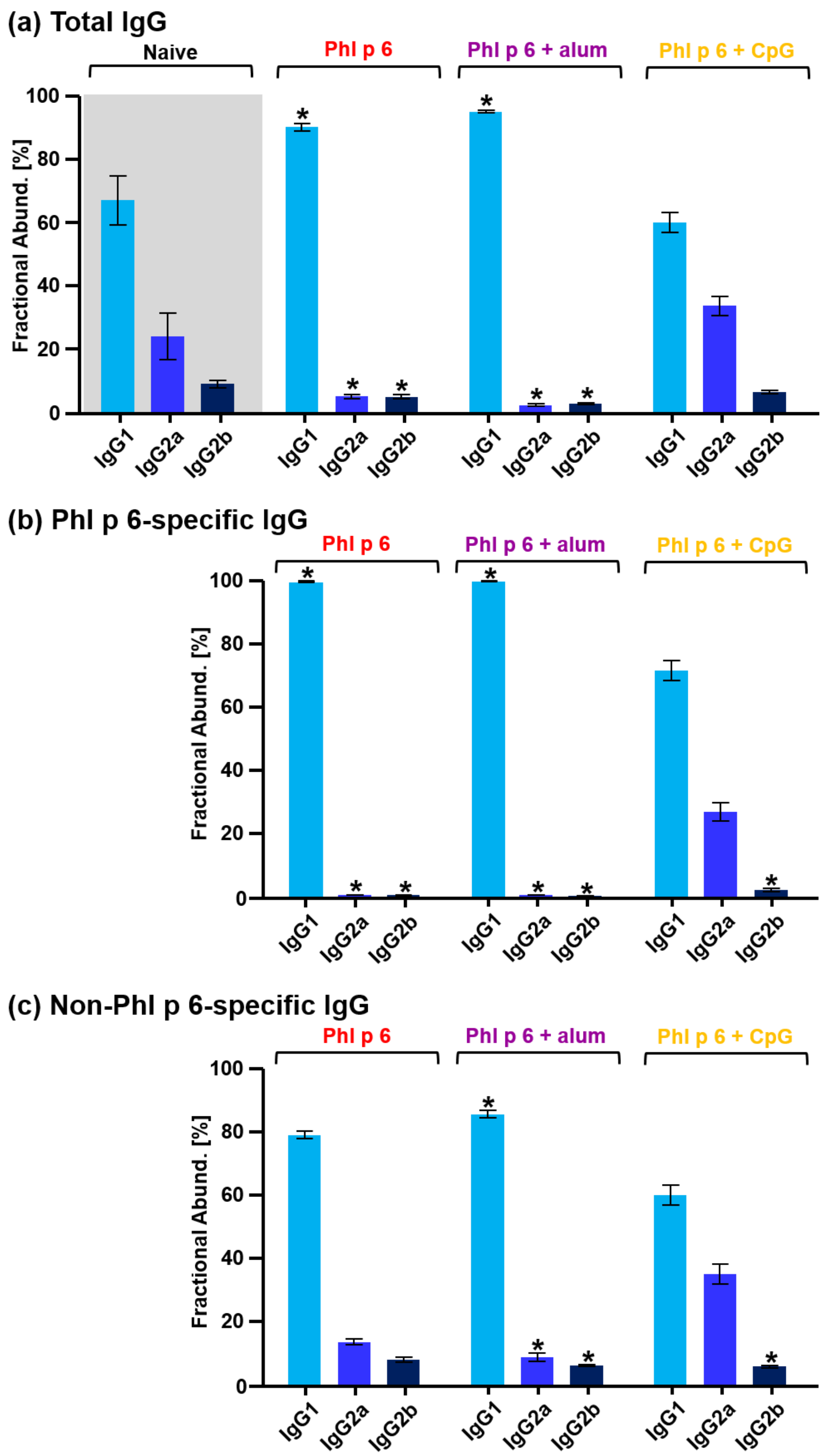


**Figure S-9.** Fractional abundances of IgG subclasses in (a) total IgG, (b) antigen-specific IgG and (c) antigen-specific depleted IgG from BALB/c mice. Abundances are shown for naive mice and cohorts vaccinated with Phl p 6, Phl p 6 + alum and Phl p 6 + CpG. Relative quantification was based on XICCs of Fc/2 subunits including all detected glycoforms. Mean values of five mouse individuals including standard deviations are shown (n=5) (Supplementary Excel file). Statistical significances were tested using a one-way ANOVA with a Dunnett post-hoc-test using naive mice as a control. Significant changes are indicated with an asterisk ($p \leq 0.05$).

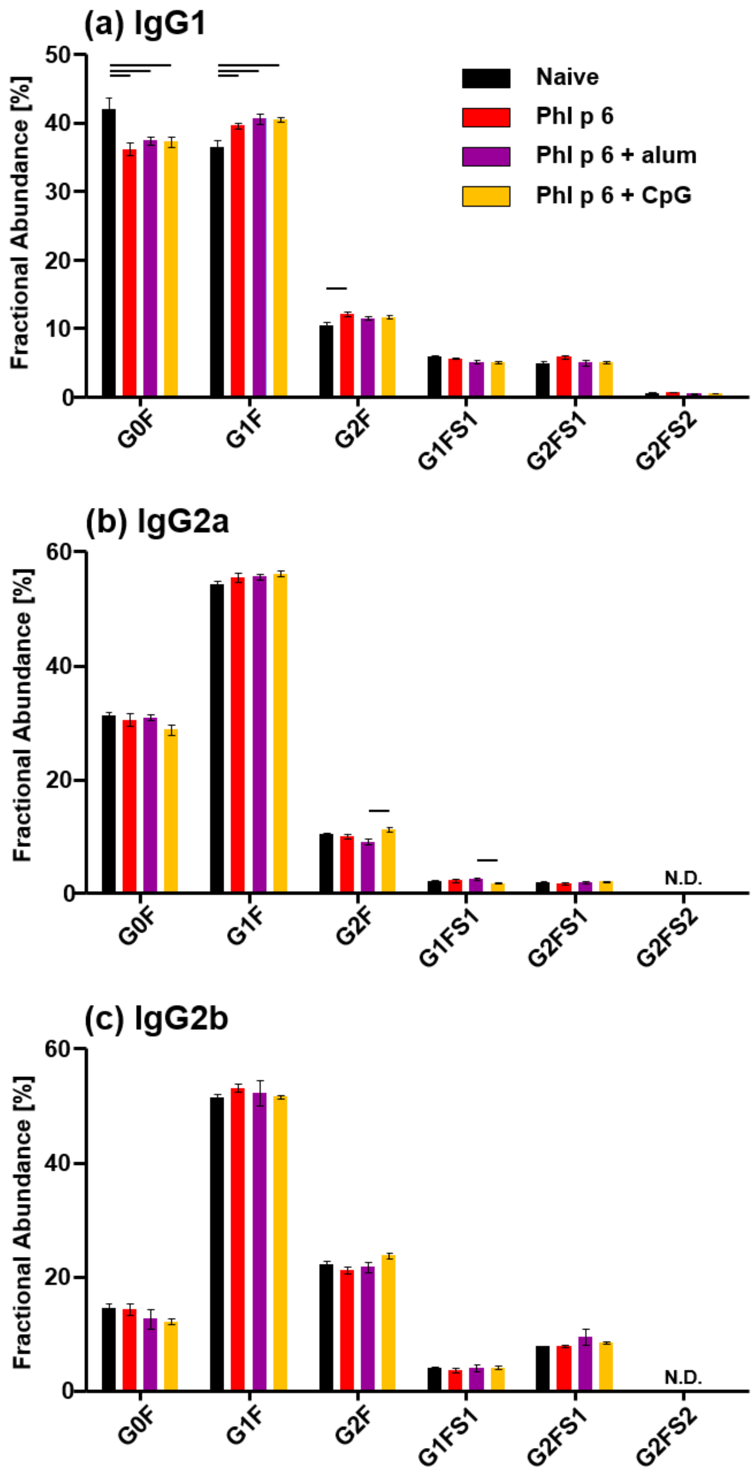


**Figure S-10.** Subclass-specific glycosylation profiles of total IgGs from naive and Phl p 6-vaccinated BALB/c mice. Relative glycoform abundances of (a) IgG1, (b) IgG2a and (c) IgG2b in naive mice and cohorts treated with Phl p 6, Phl p 6 + alum and Phl p 6 + CpG, respectively, are shown. Fractional abundances represent mean values of five mouse individuals including standard deviations (n=5) (Supplementary Excel file). Statistical significances were tested using a one-way ANOVA with a Tukey post-hoc-test. Significant changes are indicated ($p \leq 0.05$).

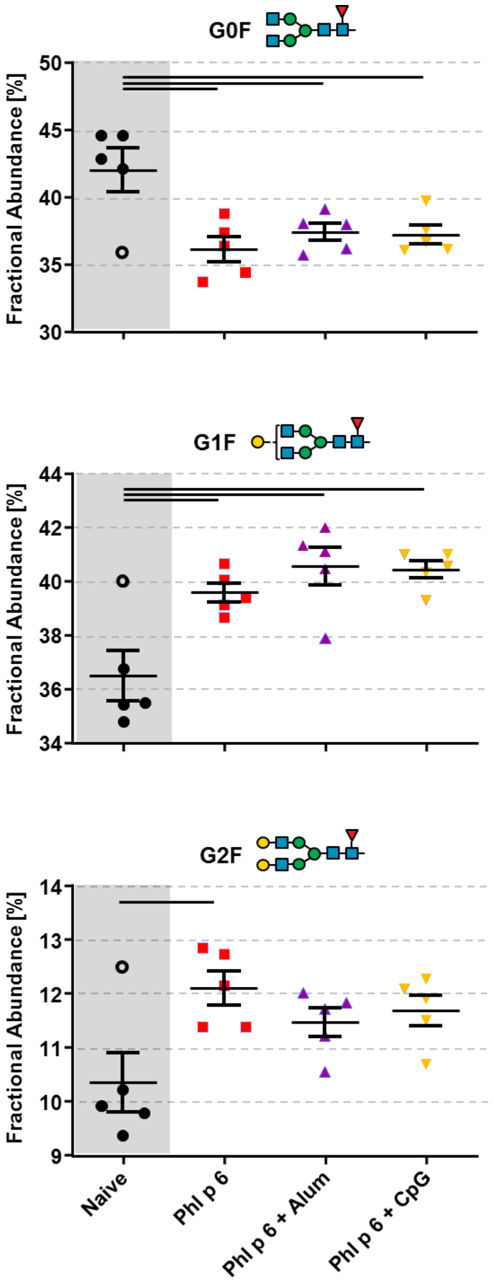


**Figure S-11.** G0F, G1F and G2F glycoform abundances of IgG1 in total IgGs from naive and Phl p 6-vaccinated BALB/c mouse individuals. Data corresponds to the bar chart shown in Figure S-10 (a). Mean values of five mouse individuals including standard deviations are shown. The outlier individual in the naive cohort is indicated as open circle. Statistical significances were tested using a one-way ANOVA with a Tukey post-hoc-test. Significant changes are indicated ($p \leq 0.05$).

**Table S-1.** SpeB cleavage products of murine IgG heavy chains from different sources. Main and minor cleavage products are designated Fc/2 and Fc/2*, respectively. Theoretical monoisotopic masses were calculated for G1F *N*-glycoforms lacking the C-terminal lysine residue; intramolecular disulphide bonds were considered to be intact. Experimental masses are mean values of three technical replicates.

[a]Theoretical monoisotopic masses were calculated using GPMAW. [b]Experimental monoisotopic masses were obtained by deconvolution using the Xtract algorithm implemented in the Xcalibur™ software. [c]Standard deviations were calculated from three technical replicates. [d]The low mass accuracy may be attributed to a previously described +1 Da shift arising from an isotopic misassignment during deconvolution.[3]

| | Subclass | Uniprot ID | Product | Amino acids | Fractional abundance [%] | Theoretical mass [Da][a] | Experimental mass [Da][b] | Standard mass deviation [mDa][c] | Mass accuracy [ppm] |
|---|---|---|---|---|---|---|---|---|---|
| BALB/c | IgG1 | P01868 | Fc/2 | 110-323 | 93.04 | 25,871.57 | 25,871.50 | 5.03 | -2.58 |
| | | | Fc/2* | 108-323 | 6.96 | 26,087.66 | 26087.87 | 15.37 | 3.96 |
| | IgG2a | P01863 | Fc/2 | 112-329 | 95.30 | 26,339.87 | 26,340.85 | 2.00 | 37.20[d] |
| | | | Fc/2* | 117-239 | 4.70 | 25,857.68 | 25,858.71 | 3.61 | 39.83[d] |
| | IgG2b | P01867 | Fc/2 | 117-334 | 64.28 | 26,088.91 | 26,089.87 | 1.53 | 36.92[d] |
| | | | Fc/2* | 116-334 | 35.72 | 26,217.00 | 26,217.01 | 4.04 | 0.25 |
| C57BL/6 | IgG1i | A0A075B5P4 | Fc/2 | 110-323 | 96.44 | 25,836.64 | 25,836.60 | 3.61 | -1.55 |
| | | | Fc/2* | 108-323 | 3.56 | 26,052.73 | 26,052.74 | 14.73 | 0.38 |
| | IgG2bi | A0A075B5P3 | Fc/2 | 118-335 | 71.69 | 26,069.97 | 26,070.98 | 1.15 | 38.87[d] |
| | | | Fc/2* | 117-335 | 28.31 | 26,198.06 | 26,198.05 | 1.00 | -0.38 |
| | IgG2c | A0A0A6YY53 | Fc/2 | 118-334 | 68.07 | 25,794.80 | 25,794.81 | 5.51 | 0.52 |
| | | | Fc/2* | 119-334 | 31.93 | 25,723.76 | 25,723.76 | 0.58 | 0.00 |
| mAb | IgG3 | P03987 | Fc/2* | 109-329 | 70.90 | 26,394.03 | 26,394.00 | 8.08 | -1.26 |
| | | | Fc/2 | 105-329 | 29.10 | 26,746.21 | 26,746.19 | 1.53 | -0.87 |

**Table S-2.** *N*-glycan structures considered in glycoform assignment using MoFi software.[4]

| Structure | Glycan | Elemental composition | Monosaccharide composition | Monoisotopic mass [Da] |
|---|---|---|---|---|
| 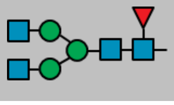 | G0F | C56H92N4O39 | HexNAc$_4$ Hex$_3$ Fuc | 1444.53387 |
| 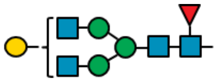 | G1F | C62H102N4O44 | HexNAc$_4$ Hex$_4$ Fuc | 1606.58669 |
| 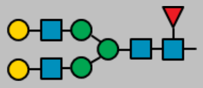 | G2F | C68H112N4O49 | HexNAc$_4$ Hex$_5$ Fuc | 1768.63952 |
| 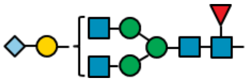 | G1FS1 | C73H119N5O53 | HexNAc$_4$ Hex$_4$ Fuc SA | 1913.67702 |
| 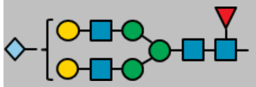 | G2FS1 | C79H129N5O58 | HexNAc$_4$ Hex$_5$ Fuc SA | 2075.72985 |
| 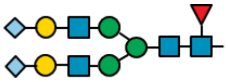 | G2FS2 | C90H148N6O67 | HexNAc$_4$ Hex$_5$ Fuc SA$_2$ | 2384.83583 |
| 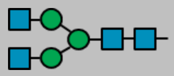 | G0 | C50H82N4O35 | HexNAc$_4$ Hex$_3$ | 1298.47596 |
| 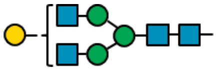 | G1 | C56H92N4O40 | HexNAc$_4$ Hex$_4$ | 1460.52878 |
| 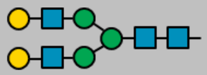 | G2 | C62H102N4O45 | HexNAc$_4$ Hex$_5$ | 1622.58161 |
| 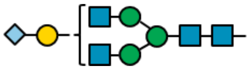 | G1S1 | C67H109N5O49 | HexNAc$_4$ Hex$_4$ SA | 1767.61911 |
| 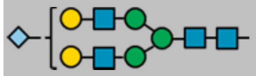 | G2S1 | C73H119N5O54 | HexNAc$_4$ Hex$_5$ SA | 1929.67194 |
| 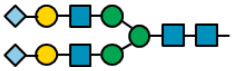 | G2S2 | C84H138N6O63 | HexNAc$_4$ Hex$_5$ SA$_2$ | 2238.77792 |
